# Seed endophytes of malting barley from different locations are shaped differently and are associated with malt quality traits

**DOI:** 10.1101/2024.05.12.593598

**Authors:** Oyeyemi Ajayi, Ramamurthy Mahalingam

## Abstract

Maximizing microbial functions for improving crop performance requires better understanding of the important drivers of plant-associated microbiomes. However, it remains unclear the forces that shapes microbial structure and assembly, and how plant seed-microbiome interactions impact grain quality. In this work, we characterized the seed endophytic microbial communities of malting barley from different geographical locations and investigated associations between bacterial species diversity and malt quality traits. Host genotype, location, and interactions (genotype x location) significantly impacted the seed endophytic microbial communities. Taxonomic composition analysis identified the most abundant genera for bacterial and fungal communities to be Bacillus (belonging to phylum Firmicutes) and Blumeria (belonging to phylum Ascomycota), respectively. We observed that a greater proportion of bacterial amplicon sequence variants (ASVs) were shared across genotypes and across locations while the greater proportion of the fungal ASVs were unique to each genotype and location. Association analysis showed a negative correlation between alpha diversity indices (Faith PD and Shannon indices) and malt quality traits for barley protein (BP), free amino nitrogen (FAN), diastatic power (DP) and alpha amylase (AA). In addition, most of the bacterial genera were significantly negatively associated with malt extract (ME) -a key trait for maltsters and brewers. We conclude that barley genotype, location, and their interactions shape the seed endophytic microbiome and is key to microbiome manipulation and management during barley production and/or malting.

## Introduction

Substantial efforts geared towards characterizing the underlying processes driving host-microbe interactions have demonstrated the indisputable role of microorganisms to agroecosystem health (French et al., 2021). The specialized symbiotic relationships between microbes and their host plants are due to their interdependence for nutrient supply (Fisher et al., 2017), and have enabled host organisms to live in habitats they would otherwise be excluded from. The microbial communities associated with plants, commonly referred to as the plant microbiome, are essential to plant growth and resistance to phytopathogens, and the seed associated microbes are co-opted early on at the seed stage to influence seedling growth vigor and plant agroecosystem adaptation (Frank et al., 2017). With seed endophytes (microbes colonizing internal seed tissues) containing specialized microorganisms driven by the host recruitment, microbiome-related traits have recently become an attractive target for breeders due to their contributions to plant health, stress tolerance and productivity. Seed endophytes like bacteria can metabolize nutrients for plant uptake (e.g. nitrogen-fixation, phosphate solubilization and siderophore production for iron uptake) and improve tolerance to abiotic and biotic stress or produce phytohormones (e.g. auxins) for plant growth and fitness (del Carmen Orozco-Mosqueda et al., 2020; Fan and Smith, 2021).

Barley (*Hordeum vulgare*) is a resilient crop that is widely distributed worldwide and is cultivated mostly in highly productive agricultural systems for food, feed, fuel, and alcohol production (Newton et al., 2011). The biodiversity of barley germplasm has led to the development of highly productive modern cultivars but also resulted in significant compositional shifts in their microbiomes (Peiffer et al., 2013; Pérez-Jaramillo et al., 2017). Yield losses due to sub-optimal cultivar performance have partly been ascribed to the significant reduction in genetic diversity of plant beneficial microbes (Doebley et al., 2006). While there may likely be an enrichment of opportunistic phytopathogens in cultivars with microbial dysbiosis, evidence highlighted the role of the host immune system in promoting microbiota homeostasis and mitigating potential virulent effects on opportunistic pathogens (Levy et al., 2017; Paasch and He, 2021).

In the field, diverse bacteria and fungi present on the barley may have originated either by vertical (acquired directly from the parent) or horizontal (acquired from the surrounding environment) transmission. In addition to barley seed carrying forward the progeny which germinates into a new plant (Nelson, 2004), they are important starting raw material in malting and brewing industry. Following harvest, microbes continue to interact with the living grain prior to malting to overcome dormancy (Bokulich and Bamforth, 2013) and the nature of their interactions could determine malt quality outcomes. For example, *Bacillus spp.* abundance can cause excessive acidification and nitrosamine formation by reducing nitrate to nitrite during wort production (Smith et al., 1992), while *Fusarium* spp, a fungal pathogen secretes mycotoxins that survive the brewing process and can be detected in finished beer (Kostelanska et al., 2009). Malting processes especially during the germination stage has been demonstrated to promote microbial activity with major influence on the malt quality downstream (Bokulich and Bamforth, 2013) and depending on the nature, number of microbes and their metabolites, the consequences for malt quality may be either deleterious or beneficial (Laitila et al., 2018). Non-viable malt derived bacteria, particularly those of submicron sizes, have been shown to disturb wort separation (Walker et al., 1997). *Pantoea agglomerans*, *Erwinia spp*, *Micrococcus spp* and *Bacillus spp.* release exopolysaccharides from the grain matrix with debilitating effects on wort filtration performance (Laitila et al., 2018). Although our knowledge of seed endophytes as beneficial microbes critical for plant growth and tolerance to biotic and abiotic stress are constantly increasing, gaps exist in our understanding of the microbial structure and diversity of the malting barley seed microbiome and their resultant effects on malt quality. To address this fundamental knowledge gap, we characterized the malting barley seed endophytic microbiomes and investigated their associations with malt quality traits.

In light of literature evidences that suggest that malt quality and wort filtration performance are significantly influenced by the growth of indigenous microbiota (Raulio et al., 2009; Bokulich and Bamforth, 2013), we asked, what are the effects of genotype (ND Genesis, Conlon and AAC Synergy) and location (St. Paul, MN; Crookston, MN; Casselton, ND and Carrington, ND) on the malting barley seed endophytic bacteria composition and assembly? Are there any associations between seed microbial taxa/diversity and malt quality traits and if so, what are the nature of these interactions? To what extent is the striking shift in microbial footprints and abundance (if any) influenced by genotype and location? We answered these questions by using next generation sequencing (16S rRNA and ITS gene sequencing) to characterize both bacterial and fungal communities, while associating bacterial diversity and abundance with malt quality.

## Materials and Methods

### Barley seed germplasm

Studies were carried out with good quality seed samples from crop year 2022 for spring two-row malting cultivar Conlon, AAC Synergy and ND Genesis. The seeds of each malting barley cultivars grown under rainfed conditions in Minnesota (St Paul and Crookston) and North Dakota (Carrington and Casselton) were pooled together to form a composite and later subsampled into two parts: the first part was set aside for bacterial and fungal endophytic characterization and the second part was set aside for micro malting.

### Surface sterilization of seed samples

One gram of barley seeds of the three malting barley varieties grown across four locations was taken in quadruplicate to give a total of 48 samples (3 cultivars X 4 locations X 4 replicates = 48 samples total) for the isolation of genomic DNA. Seed surface sterilization was carried out following a method described previously (Aswini et al., 2023). Specifically, barley seeds were first transferred to sterile 50 ml tubes and rinsed with sterile distilled water and washed with 70% ethanol for 3mins, followed by treatment with 1% sodium hypochlorite for 150 s. After sodium hypochlorite sterilization, each replicated sample was further subjected to another 70% ethanol wash and subsequently rinsed five times with sterile distilled water. To evaluate the efficiency of the sterilization, 100 µl of the water from the last rinse from each sample was plated on nutrient agar and tryptic soy agar plates and incubated at 30°C for 3 days. Sterilized seeds were subsequently dried in an oven at 50°C overnight to remove moisture and subsequently placed in minus 80°C until further analysis.

### Malt quality analyses

Malt quality analysis for each genotype-location samples were performed following methods described elsewhere (Ajayi et al., 2023). Briefly, one hundred and ten (110) grams (on a dry basis) of each barley sample were malted at the Cereal Crops Research Unit’s Malt Quality Lab (Madison, WI) to estimate malt quality parameters; diastatic power (DP), alpha amylase (AA), malt extract (ME), wort protein (WP), soluble to total protein (ST) ratio, and free amino nitrogen (FAN). The malting procedure consisted of the imbibition of barley grains (i.e., steeping) in a tank that was periodically submerged over a 36-h period to achieve a target moisture content of 45%. After reaching a 45% grain moisture content following steeping, the imbibed barley seeds were transferred to germinators and incubated under controlled temperature (16^∘^C), humidity (95%), and airflow conditions for five days. Samples were then subjected to kilning to arrest germination, preserve the modified seed, and bring the moisture levels to about 4%. After the rootlets were removed, malts were ground and subsequently analyzed for malt quality parameters following established procedures approved by the American Society of Brewing Chemists (Mahalingam, 2018).

### DNA isolation for metagenomic sequencing

For total genomic DNA, sterilized samples were grounded with liquid nitrogen and powdered seeds were processed using DNeasy Plant Mini kit (Qiagen) for genomic DNA isolation according to the manufacturer’s protocol. Following extraction, DNA quality was assessed using 0.8% agarose gel, and the purity was determined using a NanoDrop spectrophotometer (Thermo Scientific, USA). Isolated DNA was stored at −80°C for 16s rRNA and ITS amplicon sequencing.

### Library preparation and Illumina 16s and ITS sequencing

For the construction and sequencing of the v3-v4 16s metagenomic libraries, purified genomic DNA was submitted to the University of Wisconsin-Madison Biotechnology Center. DNA concentration was verified fluorometrically using either the Qubit® dsDNA HS Assay Kit or Quant-iT™ PicoGreen® dsDNA Assay Kit (ThermoFisher Scientific, Waltham, MA, USA). Samples were prepared in a similar process to the one described in Illumina’s 16s Metagenomic Sequencing Library Preparation Protocol, Part # 15044223 Rev. B (Illumina Inc., San Diego, California, USA) with the following modifications: The 16S rRNA gene V3/V4 variable region was amplified with fusion primers (forward primer 341f: 5’-ACACTCTTTCCCTACACGACGCTCTTCCGATCT(N)_0/6_<u>CCTACGGGNGGCWGCAG</u>-3’, reverse primer 805r: 5’-GTGACTGGAGTTCAGACGTGTGCTCTTCCGATCT(N)_0/6_<u>GACTACHVGGGTATCTAATC</u> <u>C</u>-3’). Region specific primers were previously described ((Klindworth et al., 2013) (underlined sequences above), and were modified to add 0 or 6 random nucleotides ((N)_0/6_) and Illumina adapter overhang nucleotide sequences 5’ of the gene-specific sequences. Following initial amplification, reactions were cleaned using a 0.7x volume of AxyPrep Mag PCR clean-up beads (Axygen Biosciences, Union City, CA). In a subsequent PCR, Illumina dual indexes and sequencing adapters were added using the following primers (forward primer: 5’-ATGATACGGCGACCACCGAGATCTACAC[5555555555]ACACTCTTTCCCTACACGAC GCTCTTCCGATCT-3’, Reverse Primer: 5’-CAAGCAGAAGACGGCATACGAGAT[7777777777]GTGACTGGAGTTCAGACGTGTGCT CTTCCGATCT −3’, where bracketed sequences are 10bp custom Unique Dual Indexes). Following PCR, reactions were cleaned using a 0.7x volume of AxyPrep Mag PCR clean-up beads (Axygen Biosciences). Quality and quantity of the finished libraries were assessed using an Agilent 4200 TapeStation DNA 1000 kit (Agilent Technologies, Santa Clara, CA) and Qubit® dsDNA HS Assay Kit (ThermoFisher Scientific), respectively. Libraries were pooled in an equimolar fashion and appropriately diluted prior to sequencing. For the construction and sequencing of the ITS libraries, samples were prepared in a similar manner to the one described in Illumina’s 16s Metagenomic Sequencing Library Preparation Protocol, Part # 15044223 Rev. B (Illumina Inc., San Diego, California, USA) with the following modifications: The ITS region was amplified with fusion primers (forward primer: 5’-ACACTCTTTCCCTACACGACGCTCTTCCGATCT<u>CTTGGTCATTTAGAGGAAGTAA</u>-3’, reverse primer: 5’-GTGACTGGAGTTCAGACGTGTGCTCTTCCGATCT<u>TCCTCCGCTTATTGATATGC</u>-3’). Region specific primers were previously described (ITS1-F (Gardes and Bruns, 1993); ITS4 (White et al., 1990)) and were modified to add Illumina adapter overhang nucleotide sequences to the region-specific sequences. Following initial amplification, reactions were cleaned using a 0.7x volume of AxyPrep Mag PCR clean-up beads (Axygen Biosciences, Union City, CA). Using the initial amplification products as template, a second PCR was performed with primers that contain Illumina dual indexes and Sequencing adapters (Forward primer: 5’-AATGATACGGCGACCACCGAGATCTACAC[55555555]ACACTCTTTCCCTACACGACG CTCTTCCGATCT-3’, Reverse Primer: 5’-CAAGCAGAAGACGGCATACGAGAT[77777777]<u>G</u>TGACTGGAGTTCAGACGTGTGCTC TTCCGATCT −3’, where bracketed sequences are equivalent to the Illumina Dual Index adapters D501-D508 and D701-D712, N716, N718-N724, N726-N729). Following PCR, reactions were cleaned using a 0.7x volume of AxyPrep Mag PCR clean-up beads (Axygen Biosciences). Quality and quantity of the finished libraries were assessed using an Agilent DNA 1000 kit (Agilent Technologies, Santa Clara, CA) and Qubit® dsDNA HS Assay Kit (ThermoFisher Scientific), respectively. Libraries were pooled in an equimolar fashion and appropriately diluted prior to sequencing. For both 16S and ITS, 300 bp paired end sequencing was performed using the Illumina NextSeq2000 as described here (White et al., 1990; Gardes and Bruns, 1993; Klindworth et al., 2013).

### Bioinformatic analysis of microbiome sequence data

Both 16S and ITS raw sequencing reads were denoised, joined, delineated into amplicon sequence variants (ASVs), and assigned taxonomy in the Qiime2 (v.2023.7) environment (Bolyen et al., 2019). For the 16S, sequence reads were demultiplexed, quality trimmed with a minimum Phred quality score of 20 (Bokulich et al., 2013) and denoised with *--p-trim-length 260* via q2-deblur in the deblur pipeline (Amir et al., 2017) to generate ASV. For the ITS, sequence reads were also demultiplexed and then filtered using the DADA2 pipeline (Callahan et al., 2016). ASVs with less than 30 reads or present in less than 3 samples within the dataset were removed to minimize potential errors in sequencing. The representative sequences were subsequently taxonomically classified using a classifier trained with the 99% OTU threshold using SILVA 138.1 database (Quast et al., 2012) for bacteria 16S, while fungal ITS sequences were classified using UNITE reference database v.9.0 (Abarenkov et al., 2024). Afterwards, qiime phylogeny align-to-tree-mafft-fasttree generated the rooted phylogenetic tree of all the ASVs for 16S sequences while reads annotated as mitochondria, chloroplasts or unassigned ASVs were culled and only reads assigned to Bacteria were considered for downstream analysis. For fungi datasets, we removed ITS reads assigned as *Viridiplantae* and *Protista* and kept only reads assigned to Fungi and both bacterial and fungal datasets were subsequently used in downstream statistical analysis.

### Statistical analyses

Downstream manipulation and analyses of the processed microbiome data generated from the bioinformatic analysis were carried out using phyloseq v.1.22.3 (McMurdie and Holmes, 2013) and Qiime2R (Bisanz, 2018). For alpha diversity, the vegan package (Oksanen et al., 2018) in R was used to perform and compute the diversity indices for both bacteria and fungi microbial communities. Kruskal–Wallis tests were performed to test for differences in alpha community diversity using QIIME2 and phyloseq in R. For beta diversity, the dissimilarity in species community composition between pairwise comparisons of bacterial and fungi communities were calculated and represented in Principal Coordinate Analysis (PCoA) ordination plots for Bray-Curtis, Unweighted Unifrac and Jaccard distances using the phyloseq package ((McMurdie and Holmes, 2013). To see if beta diversity is statistically significant between the groups, Permutational multivariate analysis of variance (PERMANOVA, 999 permutations) tests which fits linear models to distance matrices and uses permutation test was used to compute variance explained by covariates genotype, location, and genotype by location interaction effects. Core microbiome was identified using R’s microbiome package (Lahti and Shetty, 2018) using a prevalence threshold of 50% with a detection limit of 0.001, while individual ASV plots for each core microbiome was estimated using centered log ratio as described earlier (Bisanz, 2018). In addition, taxa bar plots of the top 30 most abundant phyla at appropriate taxonomic levels was generated to give an indication of how dominant phyla change between genotypes and across locations. Linear discriminant analysis (LDA) effect size which estimates magnitude of differential abundance for pairwise genotype and location comparison was conducted using lefser package in R (Segata et al., 2011) and visualized in dot plots. In addition, microbial taxa that are differentially abundant between genotypes and locations were performed using edgeR package in R (Robinson et al., 2010), which uses a TMM normalization method and negative binomial model to model raw counts of sequences observed per ASV. Using exact test in edgeR, the log2-fold change in abundance for each ASV across all pairwise contrasts was performed following adjustment for covariates. Significance was determined at p<0.05 after adjustment for multiple comparisons using the Benjamini-Hochberg false discovery rate. Venn diagrams were conducted by R package VennDiagram (v1.7.1, https://CRAN.R-project.org/package=VennDiagram). UpSet plots were generated by R package UpSetR(Conway et al., 2017), while transformed centered log ratio (CLR) used to address the compositional nature of the microbiome data was used for the ASV abundances plots and were performed using Qiime2R (Bisanz, 2018). For the association analysis involving alpha diversity metrics including bacterial genera and malt quality traits, linear mixed effect model was used to control for variation due to random effects (genotype and location) while using alpha diversity (Shannon and Faith’s phylogenetic diversity (Faith’s PD) as fixed effects. Also, correlation matrix with significance levels between alpha diversity metrics and malt quality traits was performed and visualized using R version 4.3.0 (https://CRAN.R-project.org/).

## Results

### Sequencing summary

For 16S sequencing, a total of 32,016,711 raw sequence samples were obtained. The lowest and the highest number of reads per sample were 274,889 and 747, 397 respectively with reads from other samples nestled in-between. After adaptor, primer, and chimera removal including quality and length trimming, 8,551,980 high-quality reads clustered at 99% sequence identity resulted in 1,554 ASVs. ASVs annotated as mitochondria, chloroplast or unassigned were removed, which resulted in 877 unique ASVs from 6,888,642 sequence features and were subsequently used for downstream analysis. To compare the diversity in different samples, we rarefied the data to 55,907 reads per sample and based on rarefaction curves, the sequencing depth was sufficient to capture the bacterial seed endophytic diversity (Supplementary Figure S1A). Similarly for the fungal ITS sequencing, a total of 31,110,549 raw sequence samples were observed. After adaptor, primer, and chimera removal including quality and length trimming, 25, 971,339 high-quality reads clustered at 99% sequence identity resulted in 18,110 ASVs. To compare the diversity in different samples, we rarefied the data and based on rarefaction curves, the sequencing depth was sufficient to capture the fungal microbial diversity (Supplementary Figure S1B).

### Comparative seed endophytic microbiome analysis showed significant variability in microbial community structure based on barley genotypes and locations

The predominant bacterial taxonomic composition of seed endophytes of malting barley genotypes (Conlon, ND Genesis, and AAC Synergy) across locations (Carrington, Casselton, Crookston, and St. Paul) can be seen in Figure. 1. Three major phyla, Proteobacteria, Actinobacteria, and Firmicutes were identified among the bacterial seed endophytes, with firmicutes observed to be the most abundant (Supplementary Table S1). Taxonomic classification at the genus level identified genus Bacillus as the most abundant and ranked among the top 10 genera (Figure 1A and Supplementary Figure S6). For within-location comparison across genotypes, we observed certain microbial taxa are more prevalent in certain genotypes than others. For example, Xanthomonas and Ureibacillus genera were more abundant in ND Genesis, compared to Conlon and AAC Synergy seed samples from Carrington location (Figure 1A, Supplementary Table S2). Similarly, for Casselton and St Paul locations, genus Brevibacillus was more abundant in Conlon genotype compared to AAC Synergy and ND Genesis. In addition, genus Lysinibacillus was most abundant in ND Genesis seed samples from Crookston compared to the rest of the genotype-location, while genus Xanthomonas was surprisingly more abundant for all the genotypes in Crookston compared to other genotype-location (Figure 1A, Supplementary Table S2). For the taxonomic composition of fungal communities, nine phyla were identified with phylum Ascomycota found to be the most abundant phyla (Supplementary Table S3). Taxonomic classification at the genus level identified genus Blumeria as the most abundant (Supplementary Table S4) and ranked among the top 10 genera (Figure 2A).

**Figure 1.**
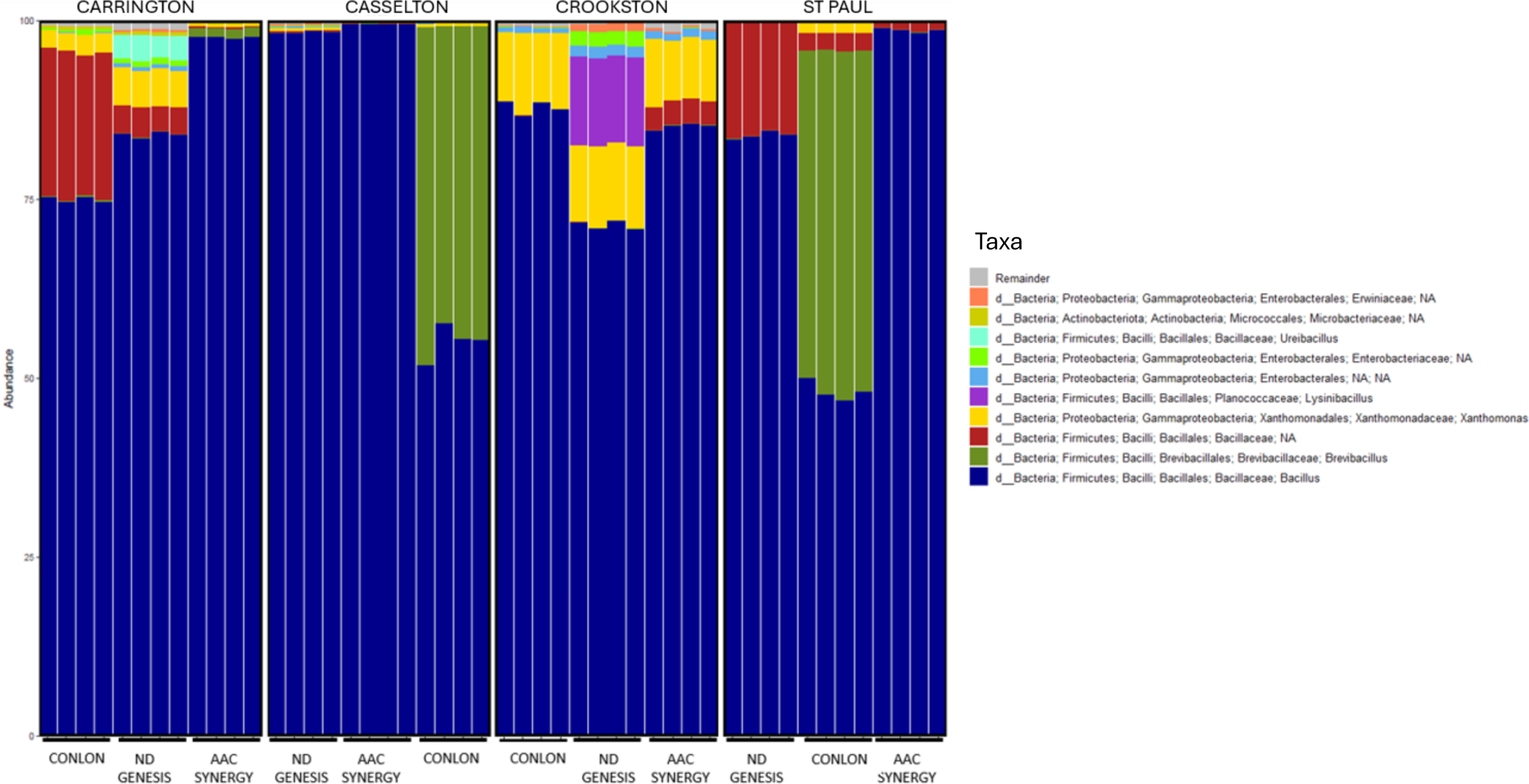
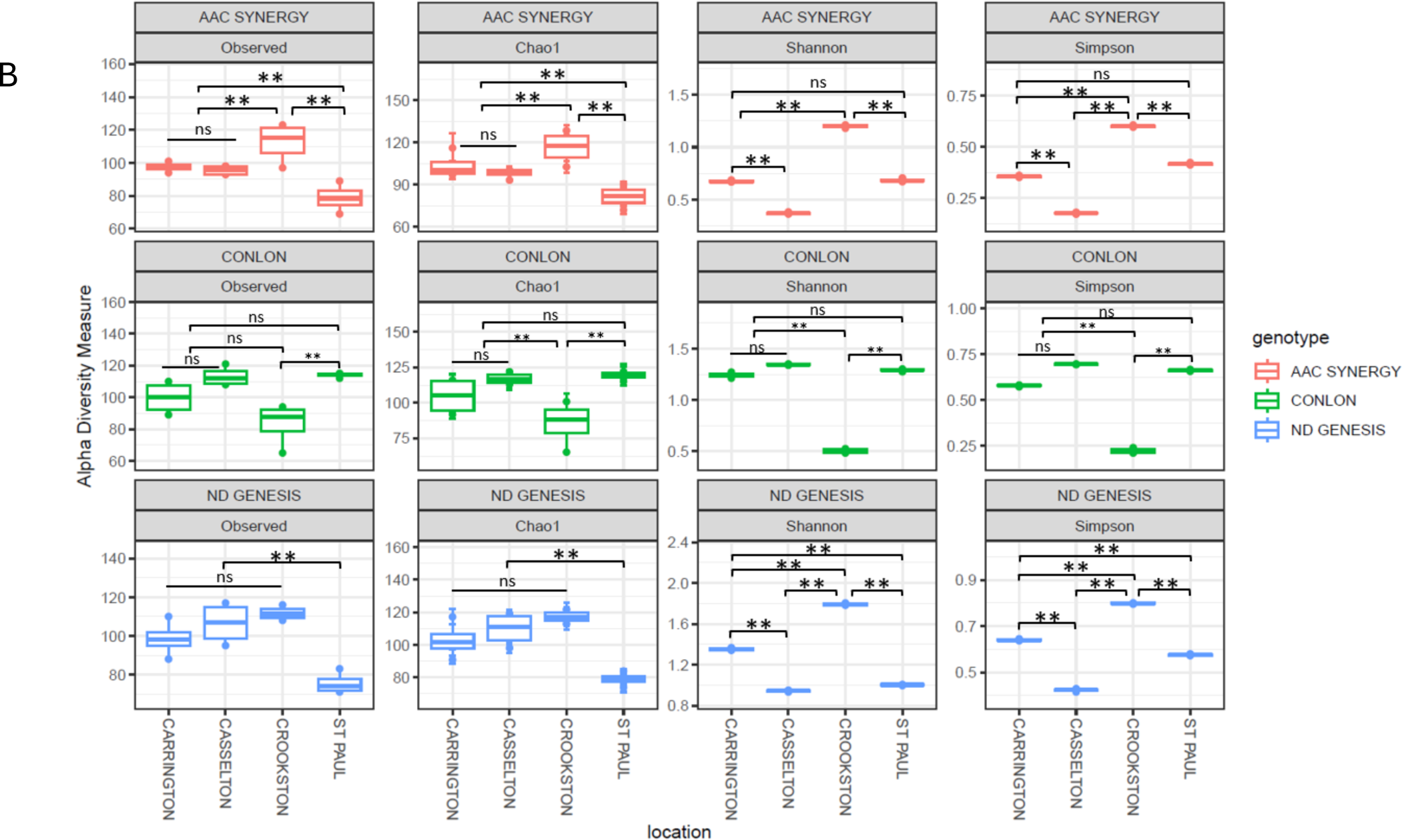
Taxonomic summary and microbial diversity analyses of bacterial seed endophytes of three malting genotype/cultivars grown across four locations. **(A)** Stacked bar chart of the taxonomic composition of bacterial communities of different genotype-location sample types aggregated at the Genus level. Each stacked column represents an independent sample (n = 48). Different colors within a column represent different genera. Only the top 10 most-abundant phyla were colored individually, all the rest are colored in gray and listed as “Remainder”. Samples are clustered by cultivar types (Conlon, ND Genesis and AAC Synergy) and location (Carrington, Casselton, Crookston and St Paul); **(B)** The bacterial seed endophytes alpha-diversity indices based on bserved species richness, Chao1, Shannon diversity index, and Simpson varied for each malting cultivar/genotype across locations (ANOVA for all the alpha diversity indices were p≤0.01). Significant asterisks ‘**\*\*’ (**p **≤** 0.01), ns = non-significant. Actual values of alpha-diversity indices can be seen in Table 1.

**Figure 2.**
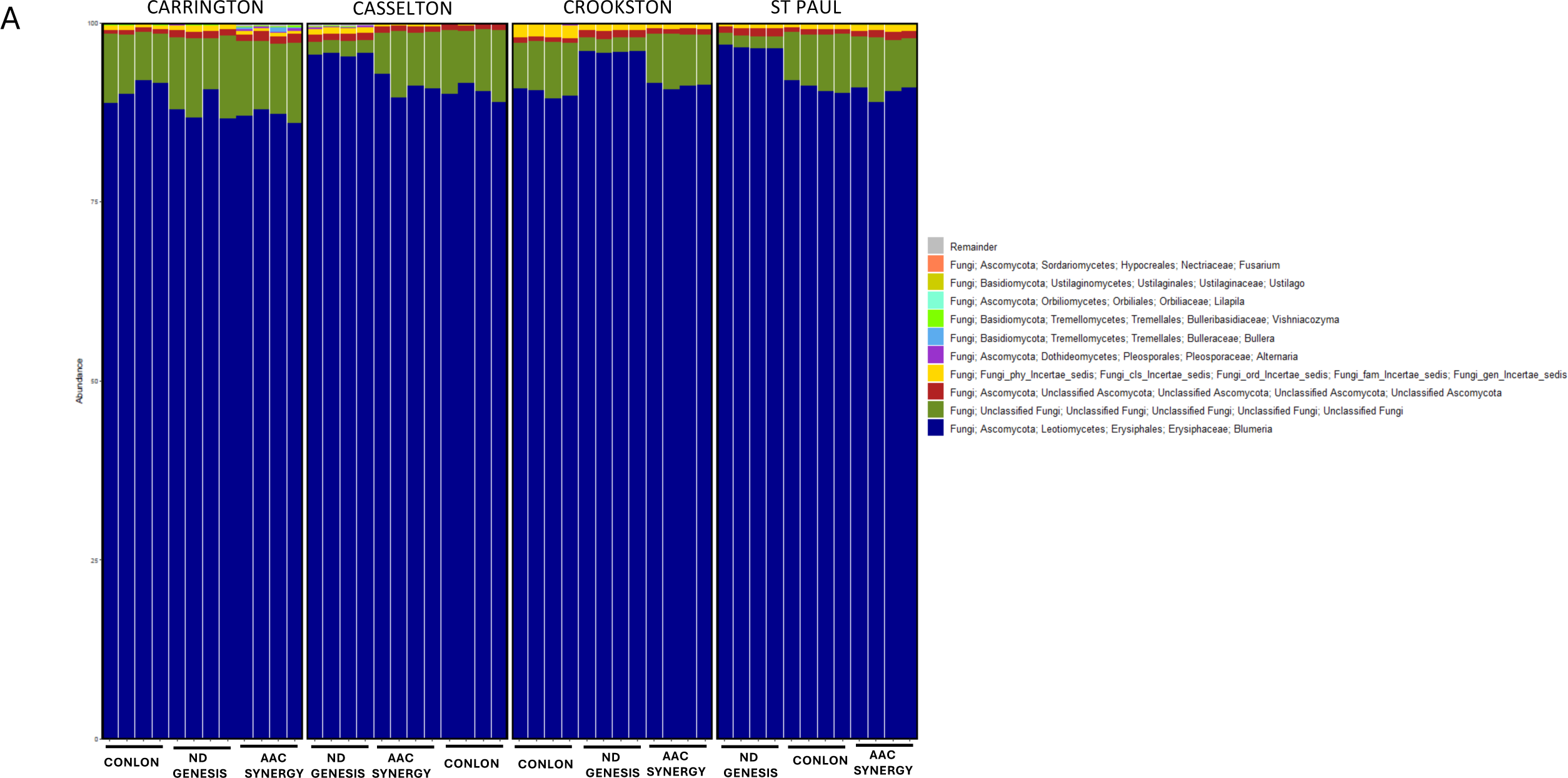
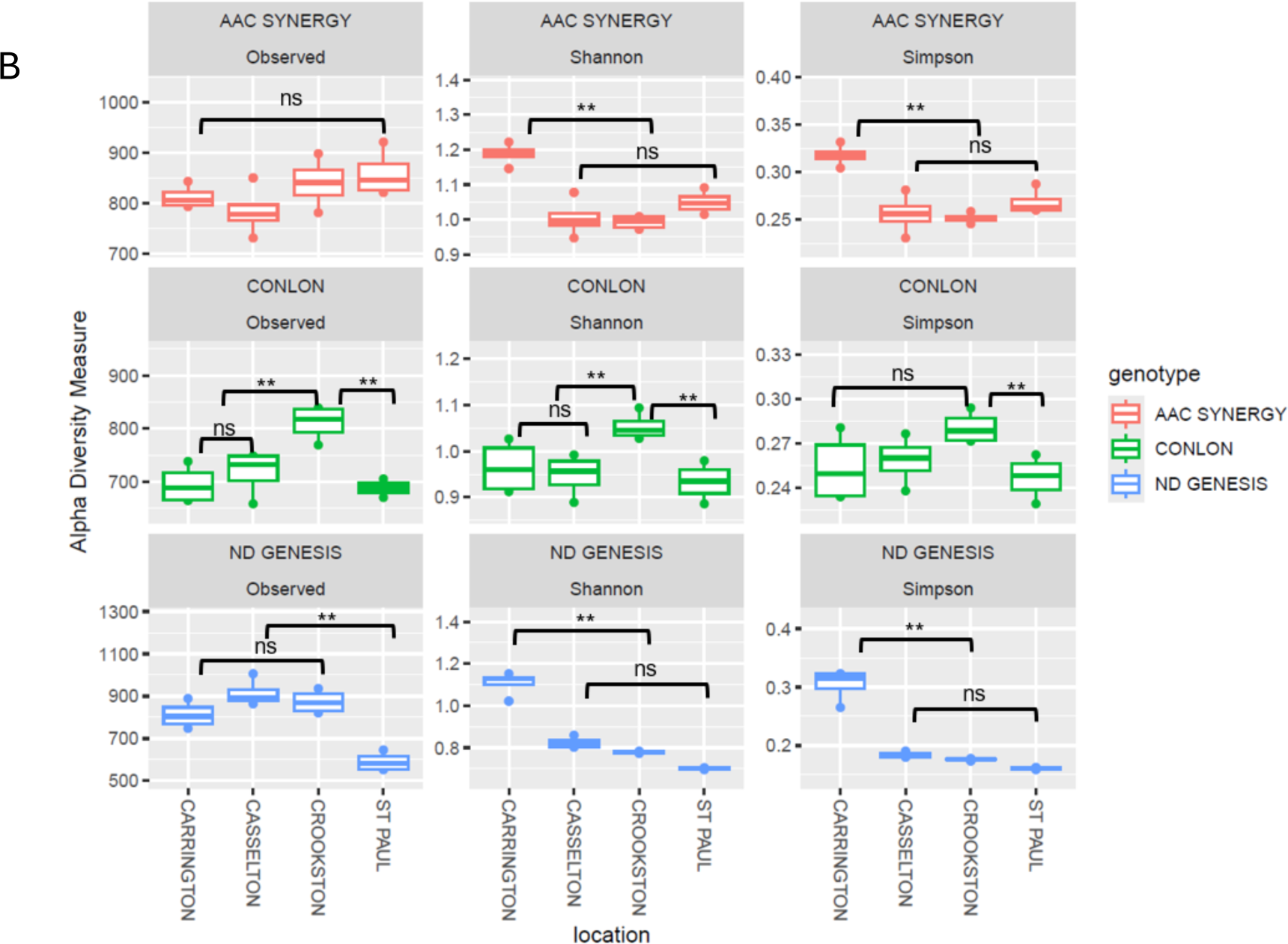
Taxonomic summary and microbial diversity analyses of fungal seed endophytes of three malting barley genotype/cultivars grown across four locations. **(A)** Stacked bar chart of the taxonomic composition of fungi communities of different genotype-location sample types aggregated at the Genus level. Each stacked column represents an independent sample (n = 48). Different colors within a column represent different genera. Only the top 10 most-abundant phyla were colored individually, all the rest are colored in gray and listed as “Remainder”. Samples are clustered by cultivar types (Conlon, ND Genesis and AAC Synergy) and location (Carrington, Casselton, Crookston and St Paul); **(B)** The fungal seed endophytes alpha-diversity indices based on observed species richness, Shannon diversity index, and Simpson varied for each malting cultivar/genotype across locations (ANOVA for all the alpha diversity indices were significant). Significant asterisks ‘**\*\*’ (**p **≤** 0.01), ns = non-significant. Actual values of alpha-diversity indices can be seen in Table 2.

**Table 1.** Alpha diversity statistics for barley seed endophytic bacterial communities (mean ± S.D)

| Genotype-location | Observed | Chao1 | Shannon | Simpson | Faith_pd |
| --- | --- | --- | --- | --- | --- |
| ND GENESIS-CASSELTON | 106.5 ± 9.23 | 109.8 ± 8.84 | 2.98 ± 0.55 | 0.42 ± 0.005 | 2.17 ± 0.12 |
| AAC SYNERGY-CASSELTON | 95.5 ± 2.5 | 97.88 ± 2.84 | 0.37 ± 0.008 | 0.18 ± 0.0002 | 1.49 ± 0.08 |
| CONLON-CASSELTON | 113.3 ± 5.21 | 117.16 ± 3.54 | 1.35 ± 0.005 | 0.7 ± 0.001 | 1.7 ± 0.19 |
| CONLON-CROOKSTON | 83.5 ± 11.46 | 85.6 ± 13.31 | 0.5 ± 0.018 | 0.22 ± 0.01 | 1.84 ± 0.09 |
| ND GENESIS-CROOKSTON | 111.8 ± 3.03 | 117.2 ± 3.45 | 1.79 ± 0.009 | 0.79 ± 0.002 | 1.83 ± 0.02 |
| AAC SYNERGY-CROOKSTON | 112.5 ± 10.43 | 116.46 ± 10.04 | 1.20 ± 0.011 | 0.6 ± 0.004 | 1.92 ± 0.08 |
| ND GENESIS-ST PAUL | 75.5 ± 4.72 | 78.87 ± 3.56 | 1.01 ± 0.003 | 0.58 ± 0.003 | 0.93 ± 0.24 |
| CONLON-ST PAUL | 114 ± 1.22 | 119.35 ± 2.66 | 1.29 ± 0.009 | 0.66 ± 0.004 | 1.54 ± 0.16 |
| AAC SYNERGY-ST PAUL | 78.75 ± 7.29 | 81.5 ± 6.89 | 0.69 ± 0.012 | 0.42 ± 0.004 | 1.19 ± 0.07 |
| CONLON-CARRINGTON | 99.75 ± 8.93 | 104.52 ± 11.07 | 1.24 ± 0.022 | 0.58 ± 0.004 | 1.8 ± 0.07 |
| ND GENESIS-CARRINGTON | 98.5 ± 7.83 | 102.82 ± 9.46 | 1.35 ± 0.010 | 0.64 ± 0.002 | 1.95 ± 0.08 |
| AAC SYNERGY-CARRINGTON | 97.5 ± 2.5 | 103.39 ± 7.6 | 0.68 ± 0.0073 | 0.36 ± 0.004 | 1.64 ± 0.07 |

**Table 2:** Alpha diversity statistics for barley seed endophytic fungal communities (mean ± S.D)

| Genotype-location | Observed | Shannon | Simpson |
| --- | --- | --- | --- |
| ND GENESIS-CASSELTON | 914 ± 54.97 | 0.82 ± 0.022 | 0.18 ± 0.004 |
| AAC SYNERGY-CASSELTON | 784.25 ± 42.54 | 1.01 ± 0.05 | 0.26 ± 0.017 |
| CONLON-CASSELTON | 717.5 ± 36.75 | 0.95 ± 0.04 | 0.26 ± 0.013 |
| CONLON-CROOKSTON | 811 ± 28.7 | 1.05 ± 0.026 | 0.28 ± 0.009 |
| ND GENESIS-CROOKSTON | 873.25 ± 47.52 | 0.78 ± 0.004 | 0.18 ± 0.001 |
| AAC SYNERGY-CROOKSTON | 840.25 ± 42.38 | 0.99 ± 0.016 | 0.25 ± 0.005 |
| ND GENESIS-ST PAUL | 590.5 ± 38.24 | 0.7 ± 0.003 | 0.16 ± 0.001 |
| CONLON-ST PAUL | 688 ± 12.98 | 0.93 ± 0.036 | 0.25 ± 0.013 |
| AAC SYNERGY-ST PAUL | 858.25 ± 39.57 | 1.05 ± 0.028 | 0.27 ± 0.011 |
| CONLON-CARRINGTON | 694.5 ± 31.12 | 0.96 ± 0.05 | 0.25 ± 0.02 |
| ND GENESIS-CARRINGTON | 811.25 ± 54.66 | 1.1 ± 0.05 | 0.30 ± 0.02 |
| AAC SYNERGY-CARRINGTON | 812 ± 19.72 | 1.19 ± 0.027 | 0.32 ± 0.01 |

Alpha diversity measures (Observed species, Chao1, Shannon and Simpson) for assessing the microbial diversity of seed endophytes based on genotype-location comparison indicated that for bacterial diversity for Observed species (number of ASVs/species when samples are rarefied) and Chao1 (measure of species richness), the highest values (114 ± 1.22 for Observed species and 119.35 ± 2.66 for Chao1 index) were found in Conlon-St. Paul while the lowest richness was detected in ND Genesis-St. Paul samples (75.5 ± 4.72 for Observed species and 78.87 ± 3.56 for Chao1 index); values for the remaining genotype-location samples are nestled between the two extremes (Table 1). Shannon and Simpson index (a metric used to estimate species diversity) and Faith’s phylogenetic diversity (Faith’s PD) which consider phylogeny to estimate diversity indicated that higher values were detected in ND Genesis-Casselton samples for Shannon and Faith’s PD (2.98 ± 0.55 for Shannon; 2.17 ± 0.12 for Faith’s PD) and ND Genesis-Crookston for Simpson index (0.79 ± 0.002 for Simpson index). Lower alpha diversity values were detected in AAC Synergy-Casselton samples for both Shannon and Simpson indices (0.37 ± 0.008 for Shannon; 0.18 ± 0.0002 for Simpson) and ND Genesis-St. Paul for Faith’s PD (0.93 ± 0.24) (Table 1). It was apparent that AAC Synergy and ND Genesis genotypes consistently showed similar patterns for all the alpha diversity measures and behaved differently from Conlon (Figure 1B). Specifically, across locations, Crookston had the highest values for the alpha diversity measured for ND Genesis and AAC Synergy and had the lowest alpha diversity values for Conlon (Figure 1B). For fungal communities, alpha diversity measures (Observed species, Shannon, and Simpson) showed that for Observed species, ND Genesis-Casselton had the highest value (914 ± 54.97) while AAC Synergy-Carrington samples had the highest values for Shannon and Simpson index (1.19 ± 0.027 for Shannon; 0.32 ± 0.01 for Simpson). Lower alpha diversity values were consistently detected in ND Genesis-St. Paul for Observed species, Shannon, and Simpson indices (Table 2). Fungal diversity across locations for AAC Synergy and ND Genesis showed similar pattern for Shannon and Simpson index except for Observed species which behaved differently (Figure 2B).

In order to investigate and visualize the seed endophytic microbiome for sample dissimilarity, we performed beta diversity measures, including Bray-Curtis distance for quantitative compositional changes (Bray-Curtis considers species abundance and species presence or absence), Jaccard distance for qualitative composition changes (Jaccard distance is based on species presence or absence) and Unweighted UniFrac distance for phylogenetic changes (Unweighted Unifrac considers presence/ absence and phylogenetic relationships). For the beta diversity of the seed bacterial endophytes microbiome, all barley genotypes from Crookston location clustered separately from the other genotype-location samples for Unweighted Unifrac, and to a certain degree for Bray-Curtis and Jaccard distances based on the two-dimension PCoA plots (Figure 3A-C; Supplementary Figure S2). For the fungal communities, distinct clustering along barley genotypes irrespective of location was observed for Jaccard distance and to a similar extent for Bray-Curtis distance (Figure 3D and Figure 3E); an exception to this was for ND Genesis where samples from Carrington clustered separately from other locations for Bray-Curtis (Supplementary Figure S3A). Permutational multivariate analysis of variance (PERMANOVA) for bacterial communities revealed significant variability among barley genotypes, with 16% in phylogeny, 35% in quantitative composition, and 27% in qualitative composition. Similarly, significant variability among barley genotypes across locations were observed with 30% in phylogeny, 26% in quantitative composition, and 27% in qualitative composition (Table 3). For fungal communities, barley genotypes showed significant variability, with 29% in quantitative composition and 27% in qualitative composition and for location, we observed a significant variability of 28% in quantitative composition and 27% in qualitative composition (Table 4). Notably, genotype and location interactions were significant for Bray-Curtis (PERMANOVA R^2^ = 0.35 for bacterial, R^2^ = 0.29 for fungal; p ≤0.001), Jaccard (PERMANOVA R^2^ = 0.29 for bacterial, R^2^ = 0.29 for fungal; p ≤ 0.001) and Unweighted Unifrac distances (PERMANOVA R^2^ = 0.28 for bacterial; p ≤ 0.001) when all samples were jointly considered for bacterial (Supplementary Table S5) and fungal communities (Supplementary Table S6). Barley genotypes, location, and their interactions significantly contributed to bacterial and fungal alpha- and beta-diversity among samples.

**Figure 3.**
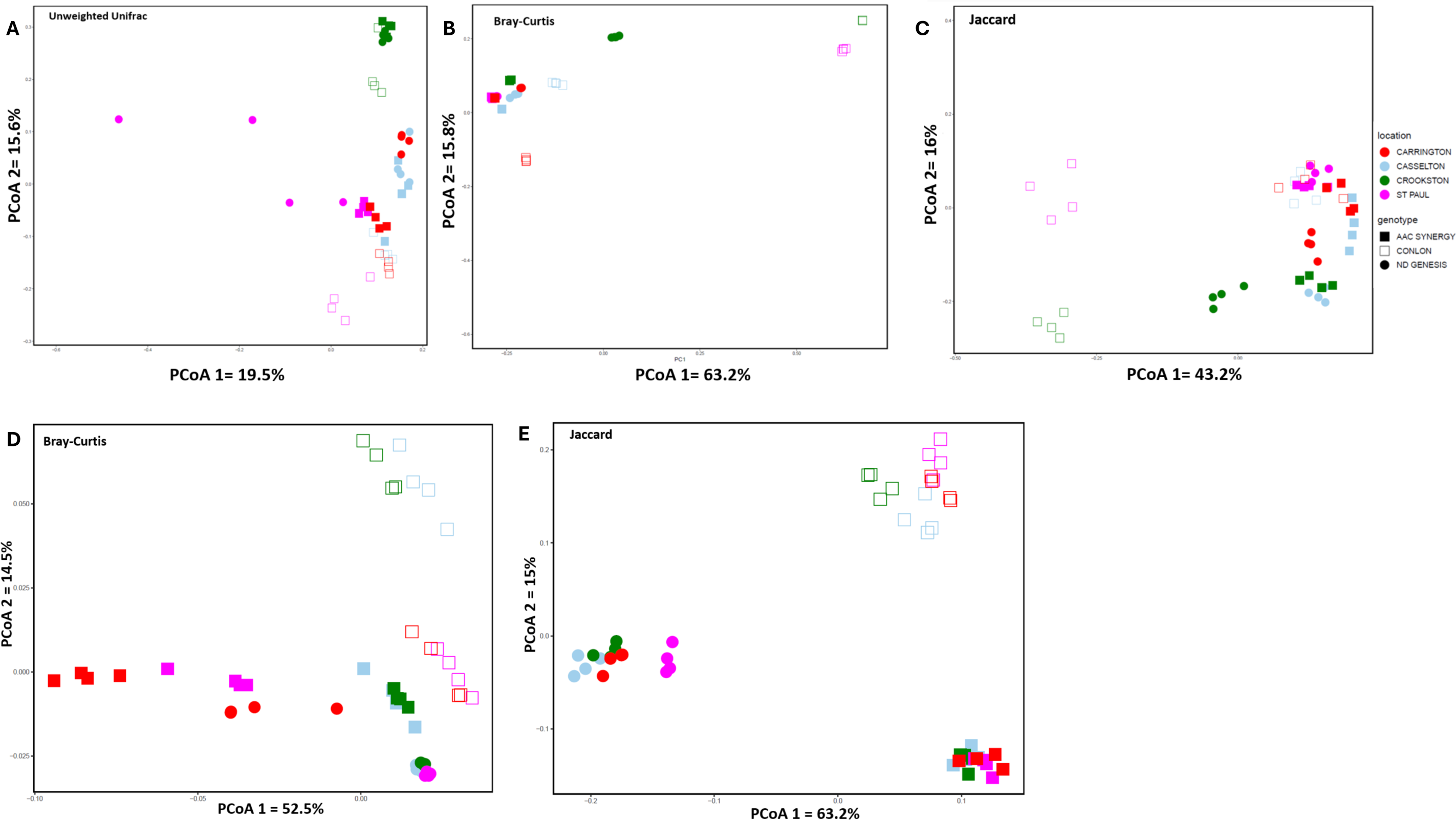
Bacterial and fungal community composition of malting barley seed endophytes grown across four locations**. A-C** represents the PCoA of 16S rRNA amplicon sequencing data while d-e represents PCoA of ITS amplicon sequencing data of genotype-location sample types based on unweighted unifrac (**A**), Bray-Curtis (**B**, **D**) and Jaccard distances (**C**, **E**). Each shape and color represents one replicate sample for genotype and location respectively for bacterial and fungal communities. Adonis in vegan package in R verified significant differences between the genotypes and locations (p≤0.001) and pairwise PERMANOVA estimates are presented in Table 3 (16s) and 4 (ITS).

**Table 3:**
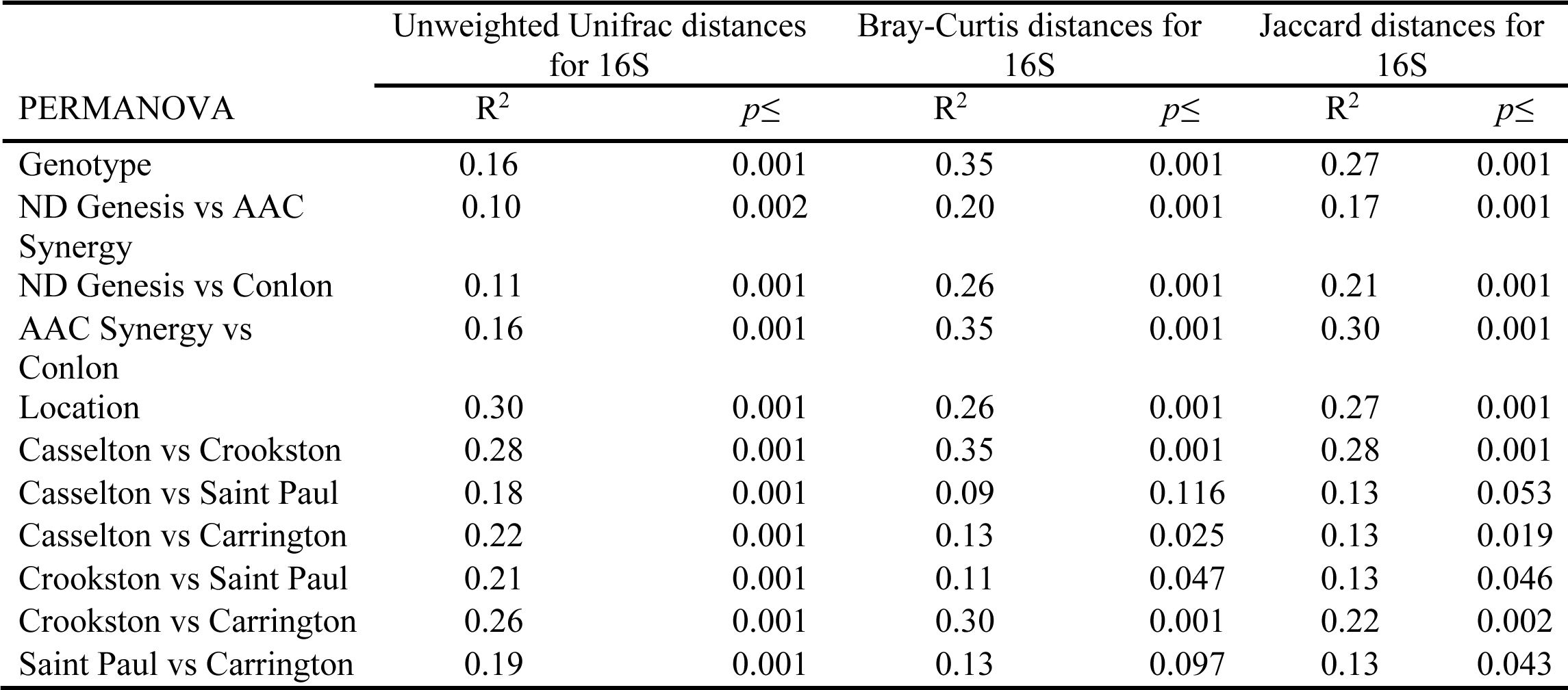
PERMANOVA of bacterial seed endophytes based on Unweighted Unifrac, Bray-Curtis and Jaccard distances.

**Table 4:**
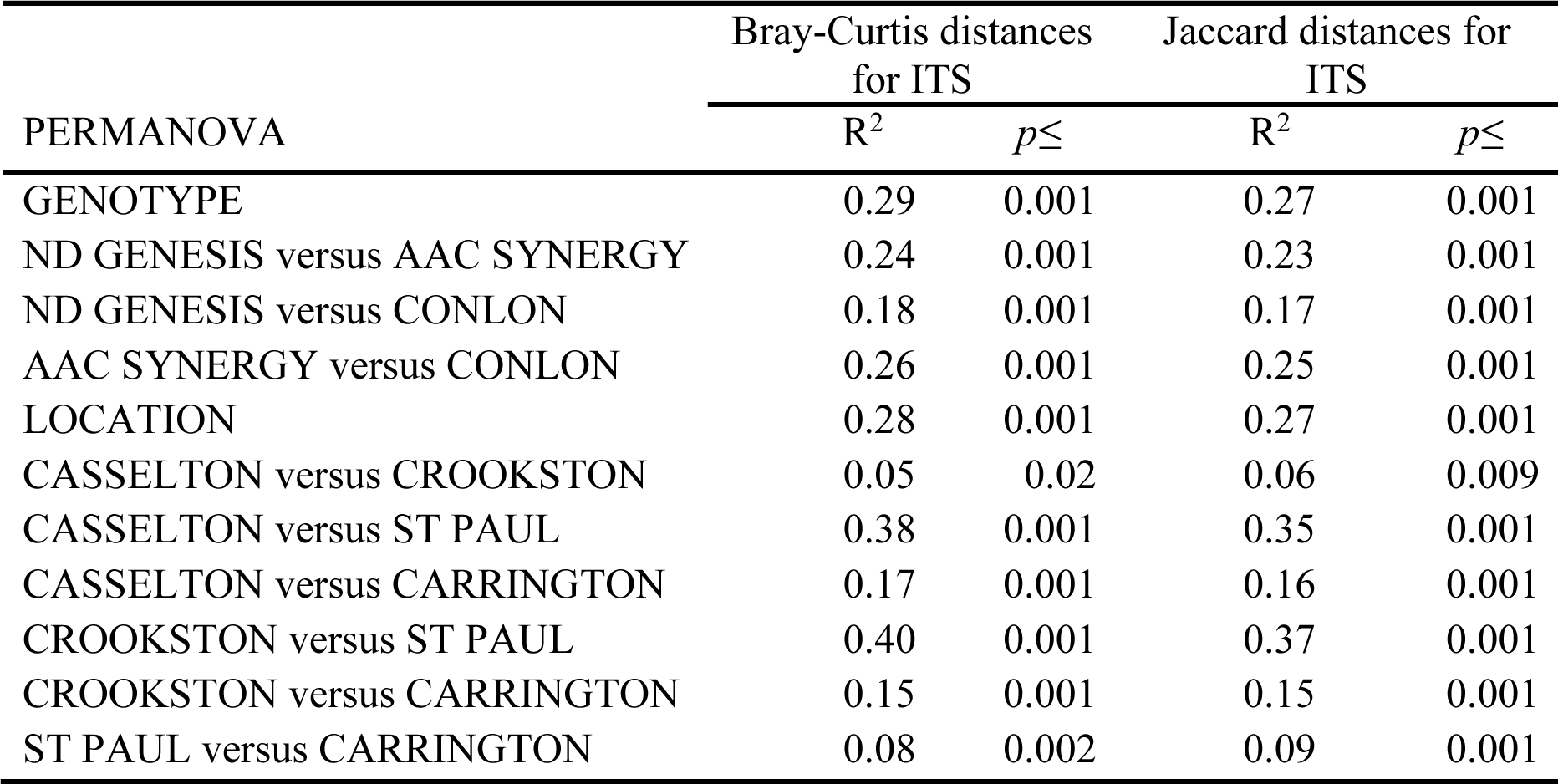
PERMANOVA of fungal seed endophytes based on Bray-Curtis and Jaccard distances for ITS.

Some alpha diversity metrics and bacterial genera are significantly associated with malt quality traits while others are not. To verify our hypothesis that bacterial genera and alpha diversity indices are associated with malt quality traits, we investigated bacterial genera and alpha diversity indices (Faith’s PD and Shannon) and their associations with malt quality traits (Figure 4). We observed that Faith’s PD was significantly negatively correlated with malt quality traits-BP (Figure 4B), FAN (Figure 4C), DP (Figure 4D), while Shannon index was significantly negatively correlated with FAN (Supplementary Figure S4A) and AA (Supplementary Figure S4B). Further analysis revealed a similar pattern of association with BP (r = −0.72; p ≤ 0.001) and DP (r = −0.78; p ≤0.001) showing the strongest correlations with Faith’s PD, and AA (r = - 0.63; p ≤ 0.001) showing moderate to high correlations with Shannon index (Figure 4F). Given the negative correlation between alpha diversity indices and malt quality, we asked whether this pattern of relationship would be detected when applied on a separate malt quality data of barley samples grown in Crookston and St. Paul, MN. We used malt quality data for crop year 2022 for St. Paul and Crookston because of the significant variability between St. Paul and Crookston for all the three genotypes (Figure 4A). Based on Welch’s t-test (due to unequal sample sizes), we found that the malt quality data for these two locations mirrors our previous findings that showed that higher alpha diversity indices (Faith’s PD and Shannon index) are negatively correlated with some malt quality traits. Interestingly, and as speculated, Crookston with significantly higher Faith’s PD had lower malt BP, DP, and FAN (Figure 4E). While AA showed significant negative relationship with Shannon index (Figure 4F), the malt quality data showed no significant difference between Crookston and St. Paul samples although Crookston AA were lower than St. Paul (Figure 4E). Notably, significantly low to moderate correlations existed between alpha diversities indices and other malt quality traits (Figure 4F). For the association between genera and malt quality traits, out of 37 genera that were identified in this study (Supplementary Table S2), only Brevibacillus (positively associated with BP), Paenibacillus (positively associated with ME), and Bacillus (positively associated with ST, WP, and FAN) had significant positive associations with malt quality traits, while the remaining genera significantly associated with malt quality traits were negatively correlated (Supplementary Figure S4C).

**Figure 4.**
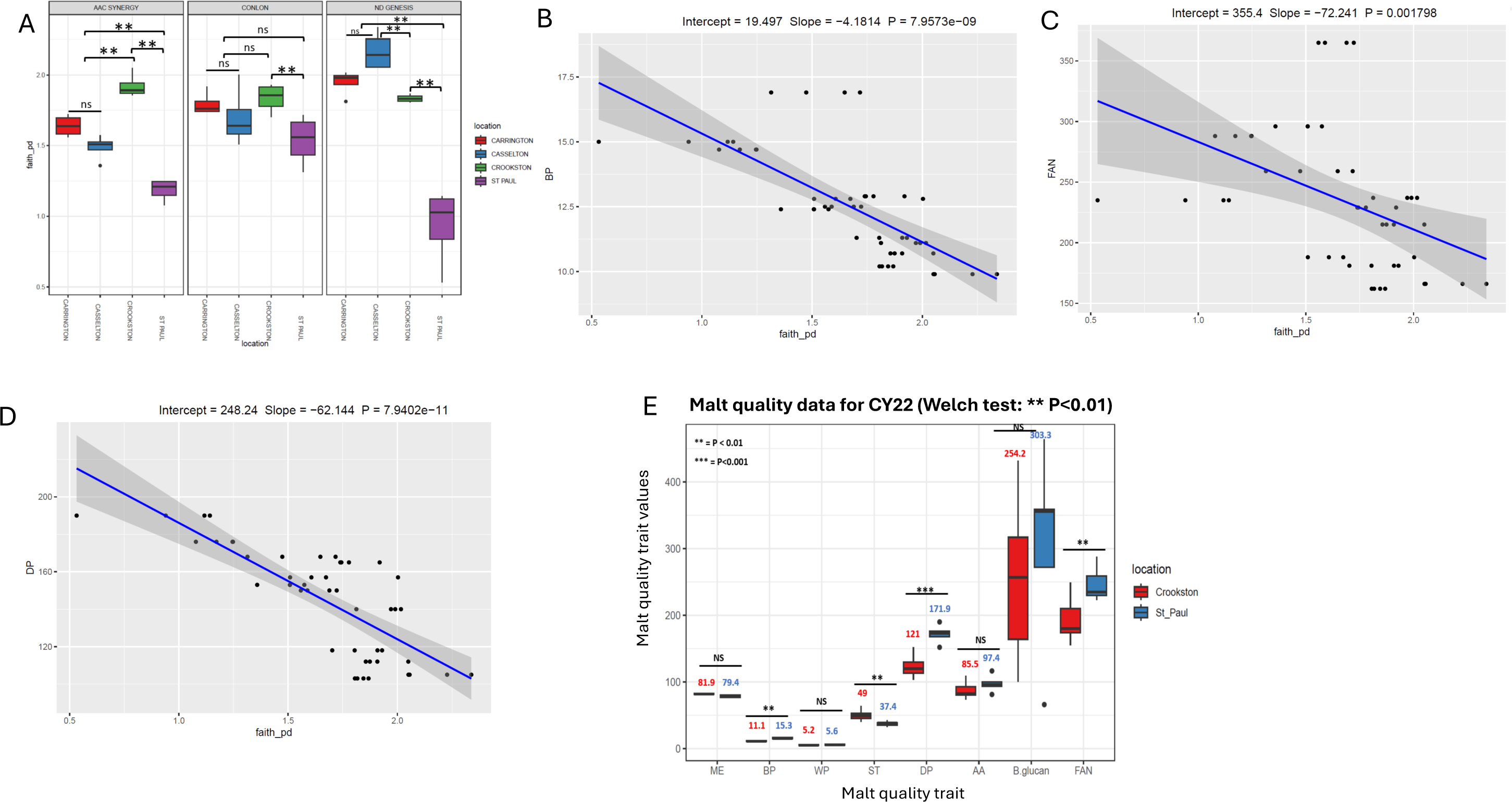
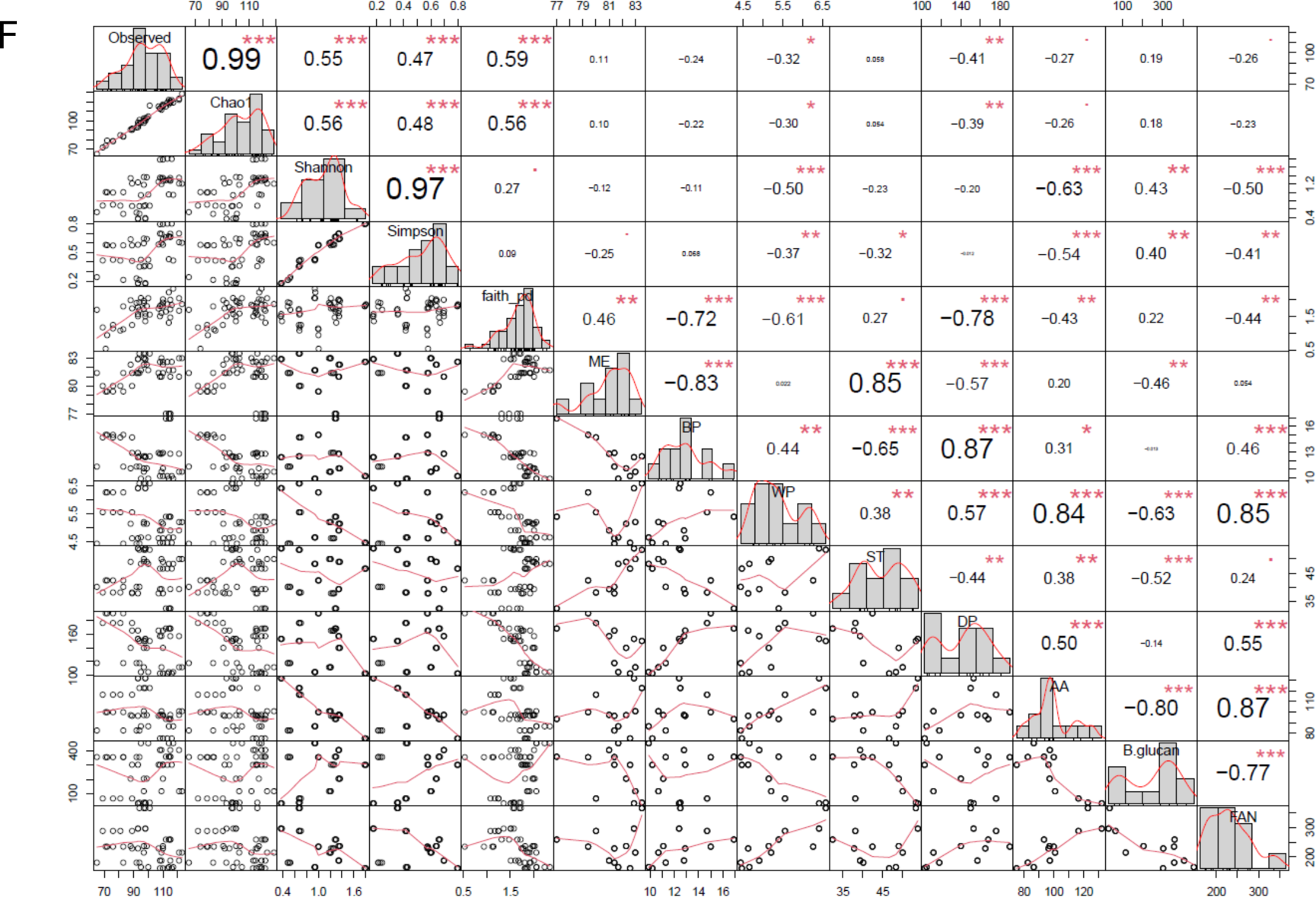
Association analysis between alpha diversity metrics, bacterial genus taxa and malt quality traits. (**A**) Alpha diversity metric (Faith_pd) of bacterial seed endophytes of three malting barley genotype/cultivars grown across four locations. (**B-D**) Negative relationship between faith_pd and BP, barley protein (**B**), free amino nitrogen (FAN) (**C**), diastatic power (DP) (**D**) based on linear regression after adjusting for covariates genotype and location; (**E**) Malt quality analyses of Conlon, ND Genesis and AAC Synergy from Crookston and St. Paul that were statistically assessed using Welch test (N = 24, 15 samples from Crookston and 9 samples from St. Paul). (**F**) Correlation analysis between alpha diversity metrics and malt quality traits. Significant asterisks ‘**\*’** indicates p **≤** 0.05; ‘**’ indicates p **≤** 0.01; ‘**\*\*\*’** indicates p **≤** 0.001.

### The core microbiome identified at the genera level and the ASVs of seed microbial endophytes differ across genotypes and locations

We defined a core microbiome as a set of microbes that are prevalent and consistently detected within a given threshold (prevalence threshold of 50% with a detection limit of 0.001) in our samples. For the bacterial core, genera Bacillus (detected in all 48 samples) and Xanthomonas (detected in 40 samples) were above the set threshold (Figure 5A and Figure 5B) and were among the top 30 bacterial seed endophytes (Supplementary Figure S6). For the fungal core, only genus Blumeria (detected in all 48 samples) was above the set threshold (Figure 5E) and the most abundant among the top 30 microbial taxa (Supplementary Figure S7). Further, BlastN analysis of bacterial genus Xanthomonas revealed that the 16s rRNA partial sequence (∼260 bp) shared 100% homology with *Xanthomonas translucens* (Supplementary Figure S8), a seed-borne phytopathogen that causes a plant disease called bacterial leaf streak. Also notable was the prevalence of Xanthomonas in all locations except ND Genesis and AAC Synergy from St. Paul location (Figure 5B).

**Figure 5.**
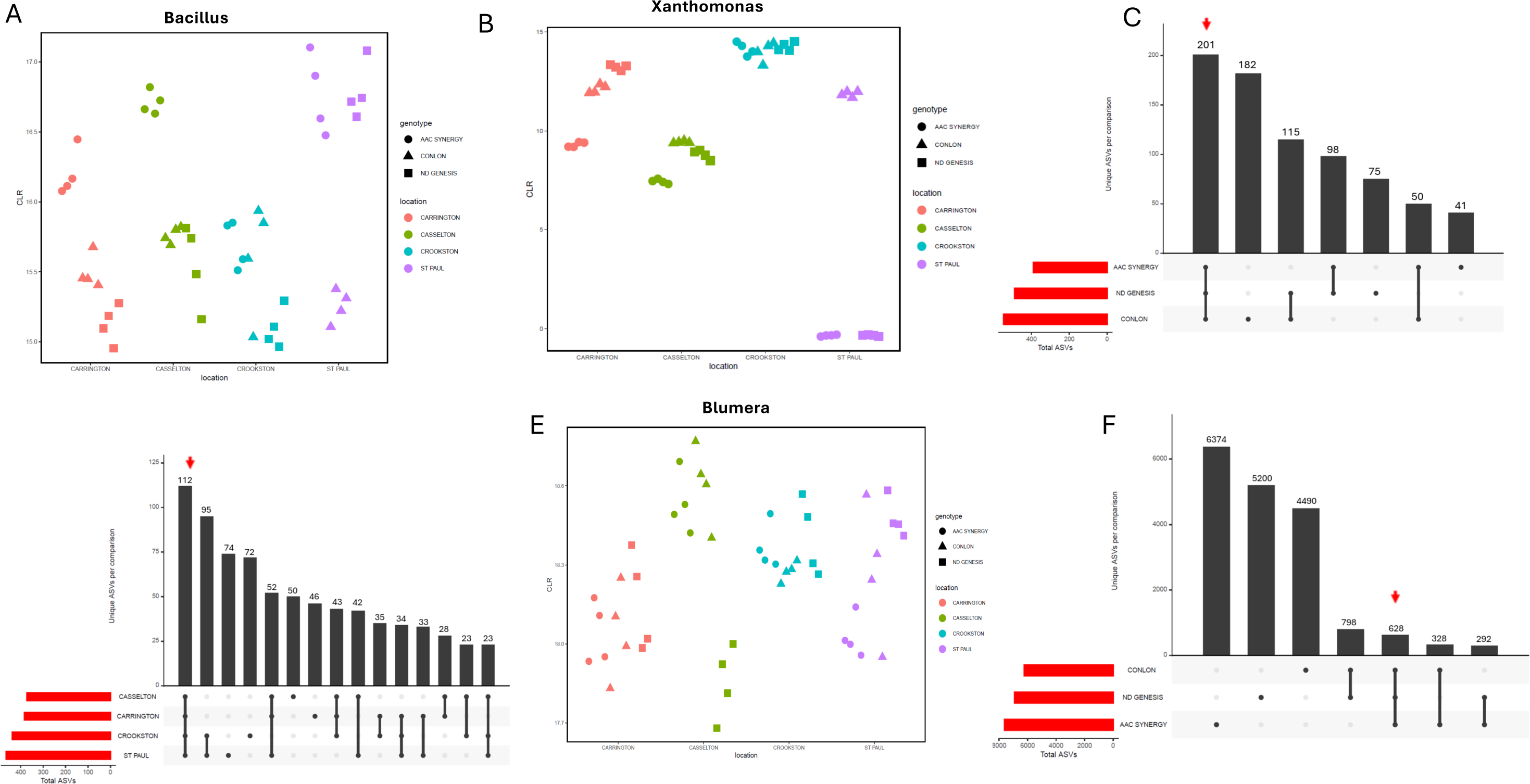
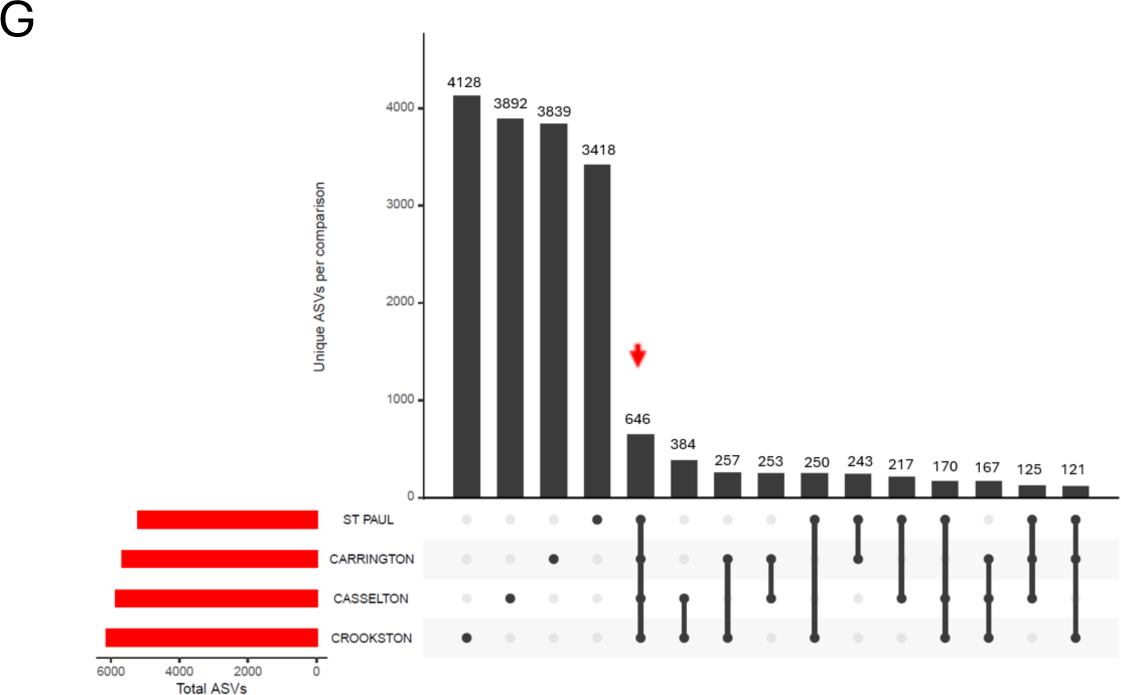
ASV abundance and UpSet plots of bacterial and fungal seed endophytes of three malting barley genotype/cultivars grown across four locations. Centered log ratio of ASV abundances for genus Bacillus (**A**) and Xanthomonas (**B**) among cultivars across location. (**C**) UpSet plot showing the number of bacterial ASVs that are shared between or are unique for AAC Synergy, ND Genesis and Conlon of all locations combined. (**D**) UpSet plot showing the number of bacterial ASVs that are shared between or are unique for Casselton, Carrington, Crookston and St Paul of all genotypes/cultivars combined. (**E**) Centered log ratio of ASV abundances for genus Blumeria among cultivars across location. (**F**) UpSet plot showing the number of fungal ASVs that are shared between or are unique for AAC Synergy, ND Genesis and Conlon of all locations combined. (**G**) UpSet plot showing the number of fungal ASVs that are shared between or are unique for Casselton, Carrington, Crookston and St Paul of all genotypes/cultivars combined. For **C,D, F, G**, red arrow indicates shared ASVs common to all locations, while filled-in black dots with an edge between the dots indicates that these ASVs are present in multiple locations.

We then focused on shared and unique bacterial and fungal ASVs between seed endophytic microbiome of all genotypes and locations. A total of 390, 489, and 548 unique bacterial ASVs were detected in AAC Synergy, ND Genesis, and Conlon seed samples, respectively. Whereas 77% ((201 + 98)/390) of the bacterial ASVs on AAC Synergy were shared with those on ND Genesis, only 65% ((201 + 115)/489) and 64% ((201 + 50)/390) of the bacterial ASVs on Conlon were shared with ND Genesis and AAC Synergy, respectively (Figure 5C). Interestingly, we found 14% (201/1427) of the bacterial ASVs present in all the three barley genotypes while 6.8% (112/1658) of the bacterial ASVs were shared across all four locations (Figure 5D). Also, we investigated core bacterial ASVs for each genotype for all locations and we observed that 1.2% (10/835), 4% (33/832) and 8.1% (63/778) of the bacterial ASVs for Conlon, ND Genesis and AAC Synergy respectively, were shared across all locations (Supplementary Figure S5A-C) indicating a higher proportion of bacterial ASVs for ND Genesis and AAC synergy is retained compared to Conlon irrespective of location. Analysis of the fungal communities showed that a total of 7622, 6918, and 6244 unique fungal ASVs were detected in AAC Synergy, ND Genesis, and Conlon seed samples, respectively. Whereas, 12% (628 + 292)/7622) of the fungal ASVs on AAC Synergy were shared with those on ND Genesis, 21% (798 + 628)/6918) and 13% (628 + 328)/7622) of the fungal ASVs on Conlon were shared with ND Genesis and AAC Synergy, respectively (Figure 5F). Only 3% (628/20,784) of the core fungal ASVs were present in all the three barley genotypes while 2.8% (646/22,818) of the fungal ASVs were shared across all four locations (Figure 5G). Also, we investigated core fungal ASVs for each genotype for all locations and observed that 4% (323/8050), 4% (346/8991) and 3.4% (326/9593) of the fungal core ASVs for Conlon, ND Genesis, and AAC Synergy respectively, were shared across all locations (Supplementary Figure S5D-F).

### Some microbial taxa are significantly more enriched or depleted in certain genotypes and locations than others

Key factors driving the observed shifts in microbiome composition identified to date include intrinsic host genotypic traits and external environmental conditions (Berg et al., 2016). Differential abundance analysis was used to investigate how genotype and location shape the overall microbial community structure (Supplementary Table S7 and S8). Pairwise comparison between St. Paul and Crookston location showed that 11 bacterial ASVs belonging to phyla Firmicutes, Actinobacteria, and Proteobacteria were significantly enriched or depleted, with Firmicutes having higher log FC for both enriched and depleted bacterial ASVs (Figure 6A). Similarly, 29 bacterial ASVs were significantly enriched for Carrington but surprisingly, no bacterial ASVs were enriched in Casselton (Supplementary Table S7), with phyla Proteobacteria, Actinobacteria, Firmicutes and Bacteroidota significantly impacted in abundance in Carrington and Casselton location (Figure 6B). Out of 11 differentially abundant bacterial ASVs for Crookston and St Paul and 29 for Casselton and Carrington, 9 were shared (Figure 6C, Supplementary Table S9). For genotype pairwise comparison, we observed 8 (Conlon and AAC Synergy; Figure 6D), 10 (ND Genesis and Conlon; Figure 6E), and 3 (ND Genesis and AAC Synergy; Figure 6F) bacterial ASVs that were significantly impacted in abundance, with more than half belonging to Firmicutes based on the three pairwise genotype comparisons (Figure 6D-F; Supplemental Table S7). While no bacterial ASVs were shared among the three pairwise genotype comparison, only 3 bacterial ASVs were shared between ND Genesis and Conlon and ND Genesis and AAC Synergy (Figure 6G). Interestingly, some of the bacterial ASVs identified in the location and genotype-based pairwise differential abundance analysis were also detected using linear discriminant analysis for investigating microbial community structure (Supplementary Figure S9). For fungal communities, 13 fungal ASVs were significantly enriched or depleted between Carrington and Casselton location and St. Paul and Crookston location (Figure 7A and Figure 7B) but only 3 fungal ASVs were shared between both location comparison (Figure 7C, Supplementary Table S9). Similarly, 16 (Conlon and AAC Synergy; Figure 7D), 21 (ND Genesis and Conlon; Figure 7E), and 9 (ND Genesis and AAC Synergy; Figure 7F) fungal ASVs were significantly enriched and depleted in abundance and were associated with phyla Ascomycota, Basidiomycota and Unclassified fungi based on the three pairwise genotype comparisons (Figure 7D, Figure 7E, and Figure 7F; Supplementary Table S8). While just one single fungal ASVs was shared among the three pairwise genotype comparison, we observed a higher number of shared fungal ASVs of pairwise genotype comparison with Conlon (11 shared fungal ASVs between Conlon and AAC Synergy and ND Genesis and Conlon) than without Conlon (Figure 7G). Interestingly, some of the fungal ASVs identified in the differential abundance analysis were similarly detected using linear discriminant analysis (Supplementary Figure S9).

**Figure 6.**
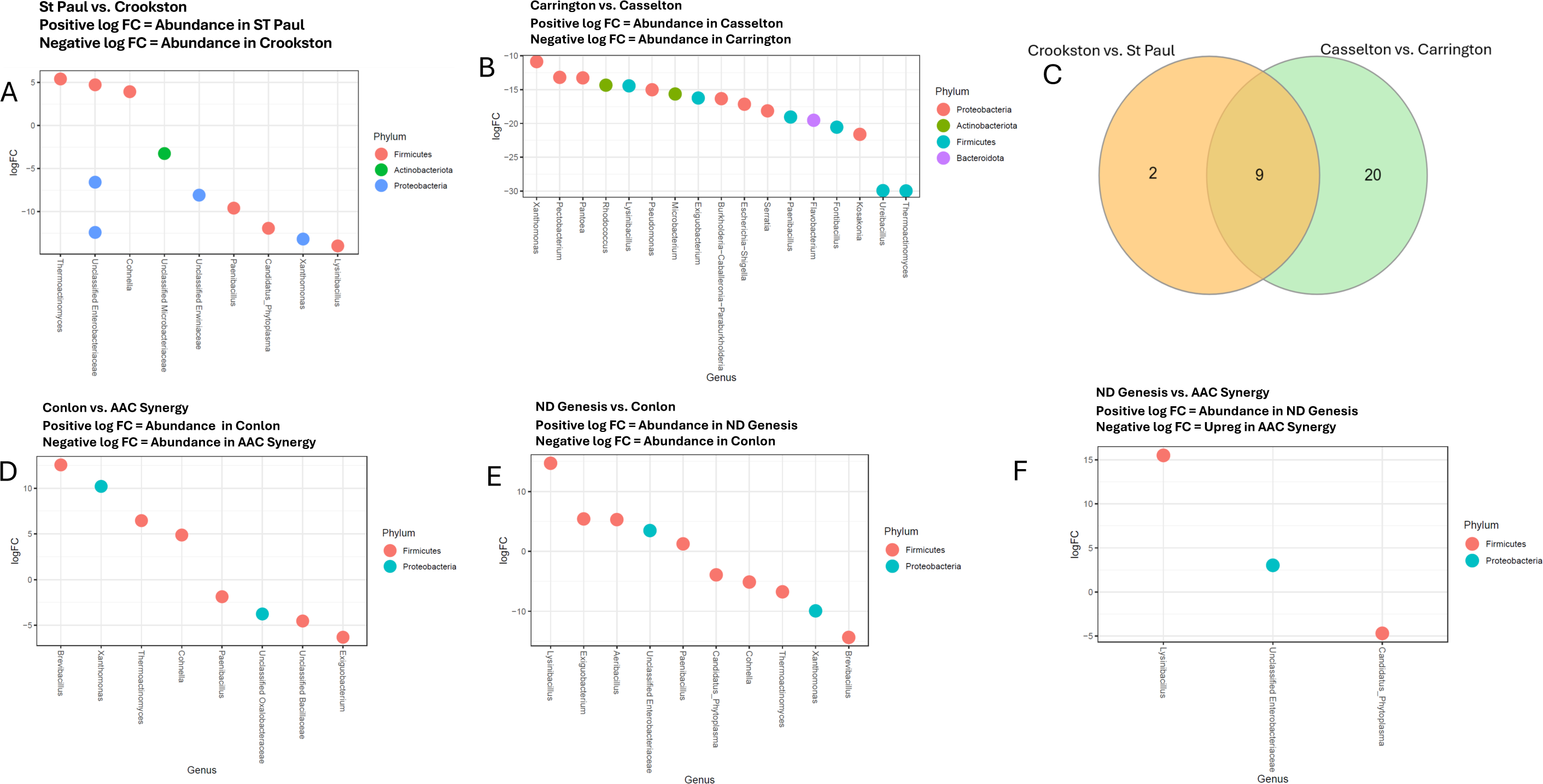
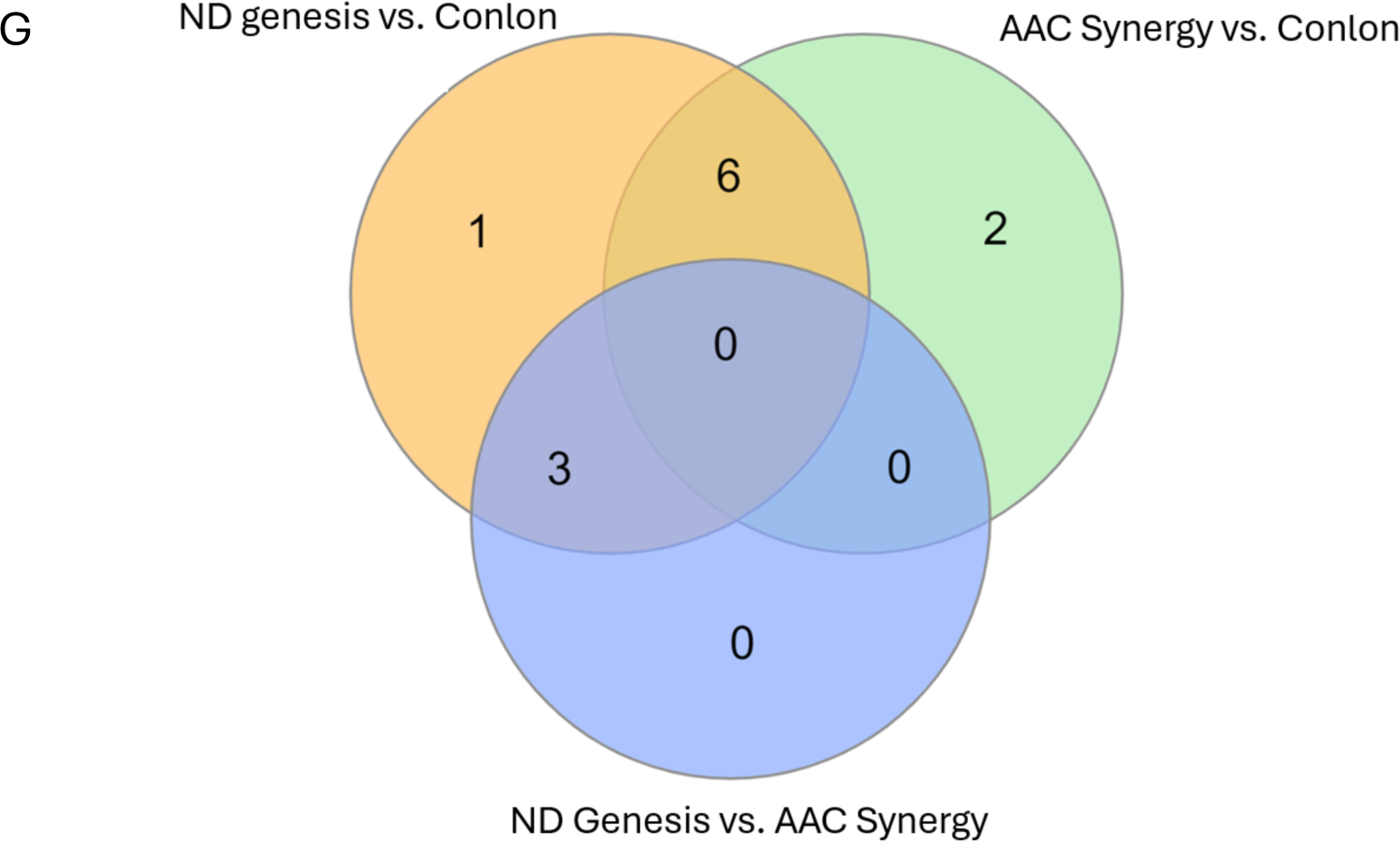
Differential abundance analysis of seed endophytic bacterial communities of malting barley grown across four locations. (**A**) Pairwise differential abundance analysis (based on logFC) between St. Paul and Crookston locations at the genus level. Positive logFC indicates increased endophytic bacterial (EB) abundance in St Paul (decrease EB in Crookston), while negative logFC indicates decreased EB in St. Paul (increased EB in Crookston). (**B**) Pairwise differential abundance analysis (based on logFC) between Carrington and Casselton locations at the genus level. The negative logFC indicates decreased EB in Casselton (increased EB in Carrington). Only some genera are reported (for complete list see Table S7) (**C**). Venn diagram showing unique and shared ASVs between pairwise location comparisons (Complete list is available in Table S7). (**D -F**) Pairwise differential abundance analysis (based on logFC) between Conlon and AAC Synergy (**D**), Conlon and ND Genesis (**E**) and AAC Synergy and ND Genesis (**F**) at the genus level. For **D**, Positive logFC indicates increased EB abundance in Conlon (decrease EB in AAC Synergy), while negative logFC indicates decreased EB in Conlon (increased EB in AAC Synergy). For **E**, Positive logFC indicates increased EB abundance in Conlon (decrease EB in ND Genesis), while negative logFC indicates decreased EB in Conlon (increased EB in ND Genesis). For **F**, Positive logFC indicates increased EB abundance in AAC Synergy (decrease EB in ND Genesis), while negative logFC indicates decreased EB in AAC Synergy (increased EB in ND Genesis). (**G**) Venn diagram showing unique and shared ASVs between pairwise genotype comparisons (Complete list is available in Table S7).

**Figure 7.**
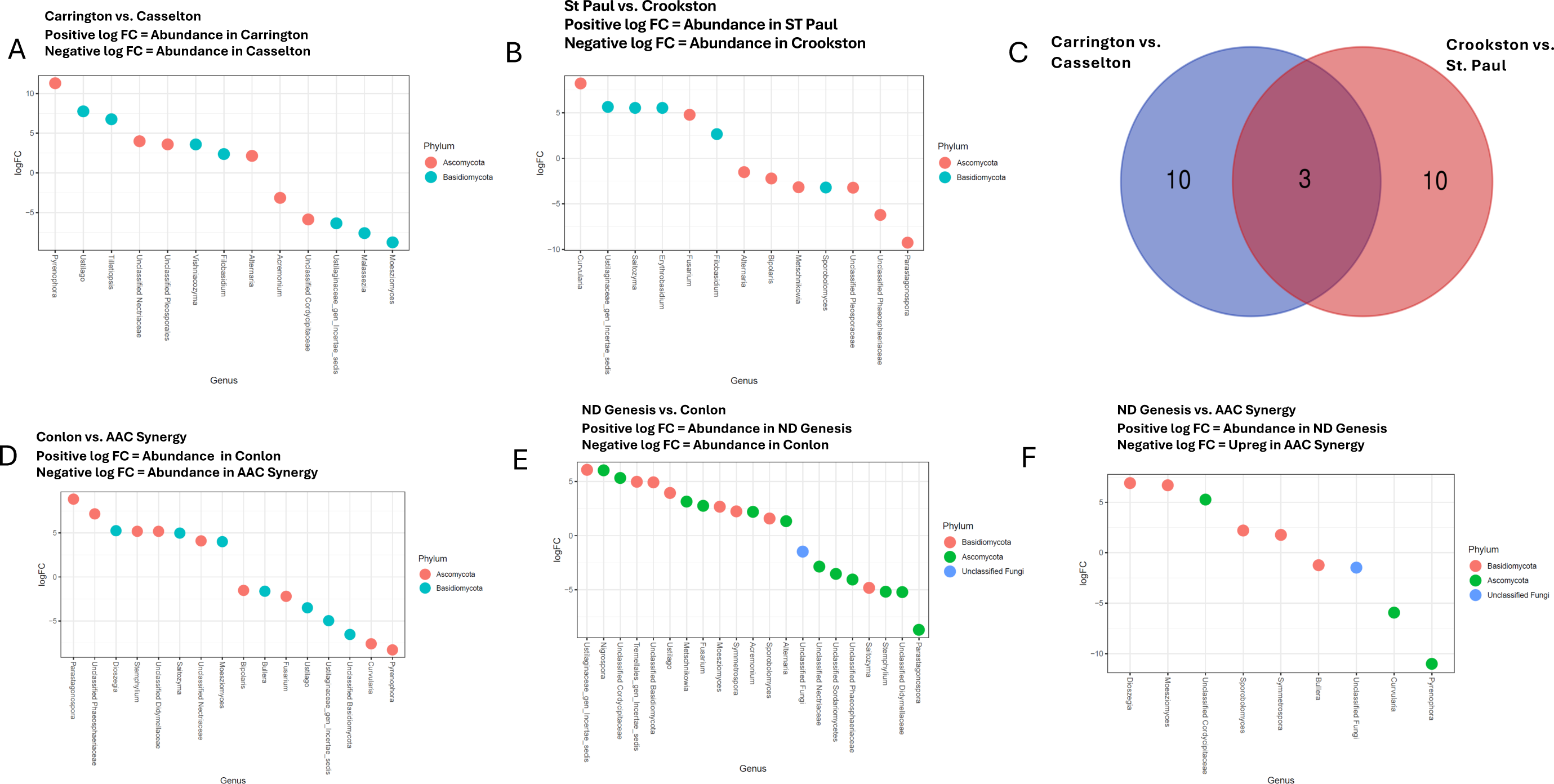
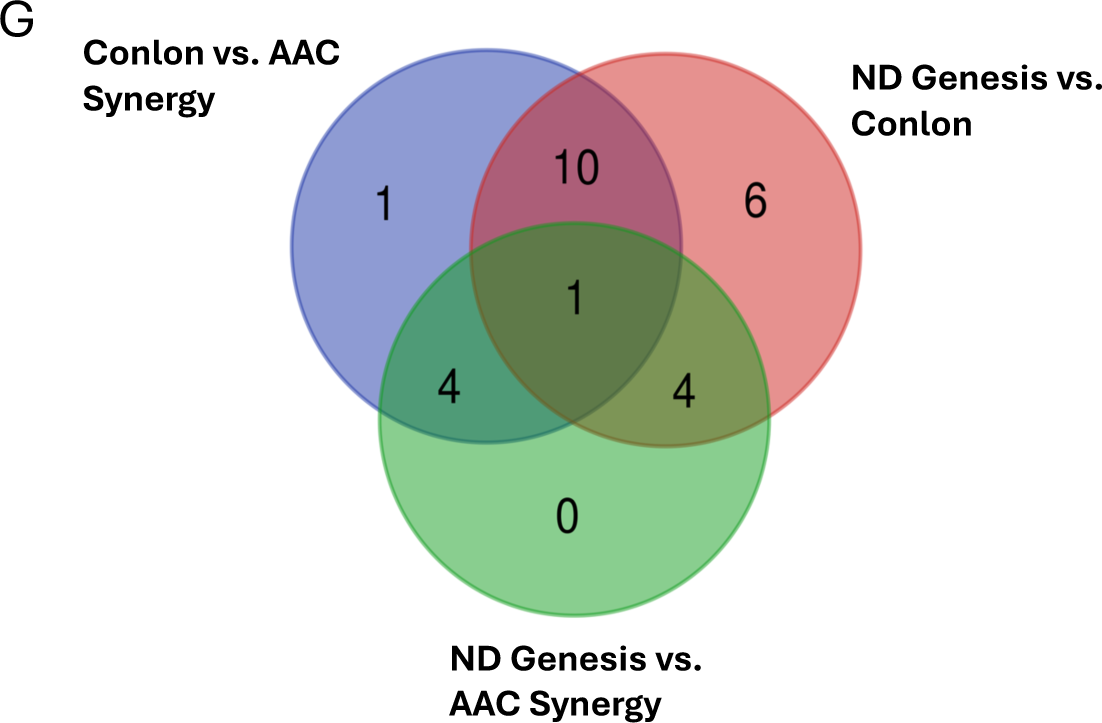
Differential abundance analysis of seed endophytic fungal communities of malting barley grown across four locations. (**A**) Pairwise differential abundance analysis (based on logFC) between Carrington and Casselton locations at the genus level. Positive logFC indicates increased endophytic bacterial (EB) abundance in Carrington (decrease EB in Casselton), while negative logFC indicates decreased EB in Carrington (increased EB in Casselton). (**B**) Pairwise differential abundance analysis (based on logFC) between St. Paul and Crookston locations at the genus level. Positive logFC indicates increased endophytic bacterial (EB) abundance in St. Paul (decrease EB in Crookston), while negative logFC indicates decreased EB in St. Paul (increased EB in Crookston). (**C**) Venn diagram showing unique and shared ASVs between pairwise location comparisons (Complete list is available in Table S8). (**D -F**) Pairwise differential abundance analysis (based on logFC) between Conlon and AAC Synergy (**D**), ND Genesis and Conlon (**E**) and ND Genesis and AAC Synergy (**F**) at the genus level. For **D**, Positive logFC indicates increased EB abundance in Conlon (decrease EB in AAC Synergy), while negative logFC indicates decreased EB in Conlon (increased EB in AAC Synergy). For **E**, Positive logFC indicates increased EB abundance in ND Genesis (decrease EB in Conlon), while negative logFC indicates decreased EB in ND Genesis (increased EB in Conlon). For **F**, Positive logFC indicates increased EB abundance in ND Genesis (decrease EB in AAC Synergy), while negative logFC indicates decreased EB in ND Genesis (increased EB in AAC Synergy). (**G**) Venn diagram showing unique and shared ASVs between pairwise genotypes comparisons (Complete list is available in Table S8).

## Discussion

Seed endophytes can be transmitted from parents to progeny and contain diverse microbial communities whose role in boosting crop yields are of particular interest for researchers. Barley seed microbiome not only perform key functions in germination, growth, biotic and abiotic stress protection (Mendes and Raaijmakers, 2015), but their reported role in influencing malt quality outcomes during the malting process has gained interest (Noots et al., 1999; Bokulich and Bamforth, 2013). Microbes are involved at every stage, from raw barley production to malting, and their functional diversity from seed exterior to interior may result either in beneficial or detrimental malt quality outcomes. To this end, we investigated the microbial community structure of malting barley seed endophytes to delineate the effects of genotype and location and the concomitant effects of their microbial composition on malt quality.

Plant genotypes has been implicated in microbial community structure, diversity, and assembly in different plant species, including *Medicago truncatula* (Brown et al., 2020), *Glycine max* (Liu et al., 2019), *Olea europaea* (Malacrinò et al., 2022), wheat (Walsh et al., 2021) and other cereals (Malacrinò et al., 2023). Our working hypothesis was that bacterial and fungal seed endophytic microbes have similar strategies for colonizing seed interior and hence face similar selection pressure, irrespective of barley seed genotypes. However, the observed results proves this not to be the case. We observed that not only genotype and location significantly influence the seed endophytic microbial communities, but their interaction effects were also significant. In fact, for some beta diversity estimates, the interaction (genotype x location) explained more variation in seed bacterial (Supplementary Table S5) and fungal microbial communities (Supplementary Table S6), than either genotype or location effects. These observations can be explained in part by the fact that seed metabolomes are unique to each genotype and their exudation may be modified by the environment to preferentially recruit certain beneficial microbes critical to plant fitness, especially when faced with onslaughts from biotic or abiotic stressors (Liu et al., 2020; Tiziani et al., 2022). In addition, the host immune system has been implicated in shaping seed endophytic microbial communities (Guttman et al., 2014; Kembel et al., 2014) and as such, host genetic control of the immune system may play a huge role in the recruitment or rejection of certain seed microbes. Alpha and beta diversity indices of barley genotypes across locations showed that AAC Synergy and ND Genesis have a similar pattern of distribution of bacterial species and abundance compared to either of them with Conlon (Figure 1B and Figure 3A-C). The very low variance in beta diversity indices (Table 3) between AAC Synergy and ND Genesis coupled with the few genera that were differentially abundant between these two genotypes (Figure 6G) suggests similarities in strategies adopted by ND Genesis and AAC Synergy in recruiting and repelling bacterial species that shaped their microbiomes. Notably, a similar pattern was observed for ND Genesis and AAC Synergy fungal communities (Figure 2B, Figure 3D and Figure 7G), albeit to a lesser extent. Also, we observed that barley samples from Crookston location consistently showed higher alpha diversity indices (Figure 1B) and clustered separately (Figure 3A, Supplementary Figure S2) from the rest of the samples. This observation further strengthens the pivotal roles of genotype, location, and their interactions in modulating seed endophytic community diversity and assembly. Notably, these observations agree with what was reported in other studies (Jung et al., 2021; Morales Moreira et al., 2021; Malacrinò et al., 2022).

The taxonomic composition analysis indicated that the most abundant genera for bacterial and fungal communities were Bacillus (belonging to phylum Firmicutes) and Blumeria (belonging to phylum Ascomycota), respectively (Supplementary Figure S6, Supplementary Table S2; Supplementary Figure S7, Supplementary Table S4). These observations were unexpected given that barley, sorghum, rice and wheat seed endophytic bacterial communities are mostly dominated by the Proteobacteria, with notable exceptions for coffee, soybeans, and *Brachypodium* seed microbiomes that are mostly dominated by the Firmicutes (Johnston-Monje et al., 2021). Bacterial species constituting the Firmicutes phylum found in C4 and CAM plants were associated with strong adaptation capacity, with Proteobacteria mostly prevalent in C3 seeds being replaced by the more resistant Firmicutes taxa in C4 and CAM photosynthetic groups (Girsowicz et al., 2019). *B. subtilis* demonstrated significant control effect on wheat powdery mildew (Xie et al., 2021) while other *Bacillus* species, such as *B. licheniformis* and *B. amyloliquefaciens* serve as biological control agents (BCAs) to control fungal diseases in the field (Cai Lin et al., 2017). Pathogen-infected plants can specifically recruit and enrich beneficial microbial communities to act as antagonists to phytopathogens (Berendsen et al., 2018). We speculate that based on these earlier observations, it appears that the Bacillus-dominated bacterial seed endophytes were specifically recruited to mitigate the effects of *Blumeria graminis*, a well-known obligate biotrophic fungus phytopathogen that causes powdery mildew impacting several economically important crops. We did not notice any discernable pattern in the abundance of Blumeria across genotypes and location except for ND Genesis in Casselton location which showed a slight decrease in the abundance based on centered log ratio estimates (Figure 5E). We observed another core genus Xanthomonas that is abundant and commonly detected in all of our genotype-location samples except for ND Genesis and AAC Synergy from St. Paul (Figure 5B, Supplemental Table S2). Xanthomonas is a major group of pathogenic bacteria infecting major cereal grains like barley, although, non-pathogenic forms have also been detected (Rana et al., 2022). To ascertain whether the genus Xanthomonas belonged to the pathogenic or non-pathogenic forms, we conducted a blastN analysis and found that the 260 bp sequence shared 100% homology with *Xanthomonas translucens* (Supplementary Figure S8), the causative pathogen for bacterial leaf streak disease (BLS) in cereals that affects grain filling. Although seeds are known to be the primary source of *X. translucens* pv. *translucens* inoculum (Duveiller et al., 1997), the prevalence and abundance of Xanthomonas as a core genus in barley seed endophytes was unexpected. We observed that ND Genesis and AAC Synergy from St. Paul were completely lacking in genus Xanthomonas and we speculate that their host immune system may play a central role in selectively repressing Xanthomonas but not *Blumeria graminis* (Figure 5B and Supplementary Table S2). Interestingly, previous work suggested the role of the host immune system in promoting microbiota homeostasis and mitigating potential invasion of opportunistic pathogens (Levy et al., 2017; Paasch and He, 2021). Considering recent evidence that indicated a reemergence of *X. translucens* pv. *translucens* within the barley-growing regions of North America (Curland et al., 2018; Beutler et al., 2023; Heiden et al., 2023; Ritzinger et al., 2023; Tambong et al., 2023), possibly due to the large-scale long-distance dissemination of the pathogen through seed (Duveiller et al., 1997), prioritizing the propagation of disease free barley genotypes in barley growing regions may be a viable option to mitigate the spread of BLS. Also worth mentioning is the appearance of a negative trend between Bacillus and Xanthomonas genera across genotype and location as the abundance of genus Xanthomonas (for example in St Paul) resulted in less abundance of genus Bacillus for Conlon genotype and this trend was noticeably consistent across our samples (Figure 5A and Figure 5B).

Seeds harbor microbiota that can be transmitted to the plants that develop from them (Berg and Raaijmakers, 2018) and understanding how microbial diversity changes across genotypes and locations could be critical to plant fitness. In this study, we observed that a greater proportion of bacterial ASVs were shared across genotypes (Figure 5C) and across location (Figure 5D) while the greater proportion of the fungal ASVs were unique to each genotype (Figure 5F) and location (Figure 5G), indicating that the underlying mechanism that shapes the recruitment of bacterial and fungal seed endophytes differ potentially due to differences in selective pressure.

The microbiological status of the grain pre- and postharvest, and during different stages of malting could influence malt quality outcomes (Bokulich and Bamforth, 2013). aAssociation studies investigating the link between human gut microbiome and human personality traits (Johnson, 2020), obesity (Murugesan et al., 2018), asthma (Attar, 2015), as well as mental health conditions (Adesman et al., 2017) have inspired us to investigate associations between plant microbial communities and economically important traits. In this study, association analysis suggests a negative correlation between alpha diversity indices (Faith’s PD and Shannon indices) and malt quality traits for barley protein (BP), free amino nitrogen (FAN), diastatic power (DP) and alpha amylase (AA) (Figure 4B-D, Figure 4G and Supplementary Figure S4A and S4B). Given these observations, we investigated whether the nature of the relationship between Faith’s PD of St. Paul and Crookston location (Figure 4A) would replicate the negative relationship between malt quality traits and Faith’s PD (for example, increase in Faith PD in Crookston would be associated with a decrease in BP, FAN, DP, and AA) when applied to a different malt quality dataset from these two locations. To our surprise, we observed that the same pattern of negative correlation between Faith’s PD and some malt quality traits (Figure 4F) were detected with significantly higher values observed for BP, DP, and FAN for St Paul samples (lower Faith’s PD) and significantly lower values for BP, DP, and FAN for Crookston samples (higher Faith’s PD). Based on these findings, we argue that increased bacterial abundance negatively influence malt quality traits; an idea consistent with previous findings that suggested that excessive bacterial growth on barley grain could retard mash filtrations due to increased abundance of bacterial exopolysaccharides (Haikara and Home, 1991; Laitila et al., 2018) and due to microbial interference with respiration during the malting process (Rani and Bhardwaj, 2021). Malt extract is a key malt quality trait as malts with high extract are desired in the brewhouses. Bacterial genera like Sphingomonas produce gellan using starch as a substrate (Freitas et al., 2011; Nwodo et al., 2012) while Pseudomonas are known to produce a variety of different bacterial polysaccharides (Bajaj et al., 2006; Vu et al., 2009). Members of genus Pantoea are well known for the production of exopolysaccharides and have been implicated in reducing wort filtration performance (Laitila et al., 2018). In this study, we observed that most of the bacterial genera were significantly negatively associated with malt extract except Paenibacillus that was significantly positively associated with malt extract (Supplementary Figure S4). This observation may be partly due to the ability of Paenibacillus to produce significant amount of xylanase compared to other bacteria in malting environment (Malfliet et al., 2013), with increased xylanase levels negatively correlated with wort viscosity and increased extract yield and brewing filterability (Cornaggia et al., 2019). Notably, our work demonstrated the significant abundance of Paenibacillus in certain genotypes and locations at the exclusion of others (for example, Paenibacillus is only present in Crookston and not St. Paul for all the genotypes; Supplementary Table S2 and Supplementary Table S7) and may influence malt quality traits. Considering these observations, future work aimed at investigating the microbial shifts (abundance and depletion) across different malting stages and their influence on malt quality traits is warranted.

## Data availability statement

The bacterial and fungal sequence data generated in this study using NextSeq2000 have been deposited and are available in the NCBI Sequence Read Archive (SRA) under Bioproject PRJNA1108745.

## Author contributions

OA: Investigation, Writing - original draft, reviewing and editing, Conceptualization, Conducted all the bioinformatics and statistics; RM – funding, Writing - review and editing.

## Acknowledgement

Authors thank Dr. Kevin Smith (University of Minnesota) and Dr. Richard Horsely for providing sample materials and metadata associated with this work. We will also want to thank the director of USDA malt quality lab, Dr. Jason Walling and his staff for providing the data for the malt quality analysis. The author (s) utilized the University of Wisconsin – Madison Biotechnology Center’s DNA Sequencing Facility (Research Resource Identifier – RRID:SCR_017759) to conduct the 16S rRNA and ITS sequencing and would like to thank them for their service. Mention of trade names or commercial products in this publication is solely for the purpose of providing specific information and does not imply recommendation or endorsement by the U.S. Department of Agriculture. USDA is an equal opportunity provider and employer.

