## Supplementary Figure for "Seed endophytes of malting barley from different locations are shaped differently and are associated with malt quality traits"

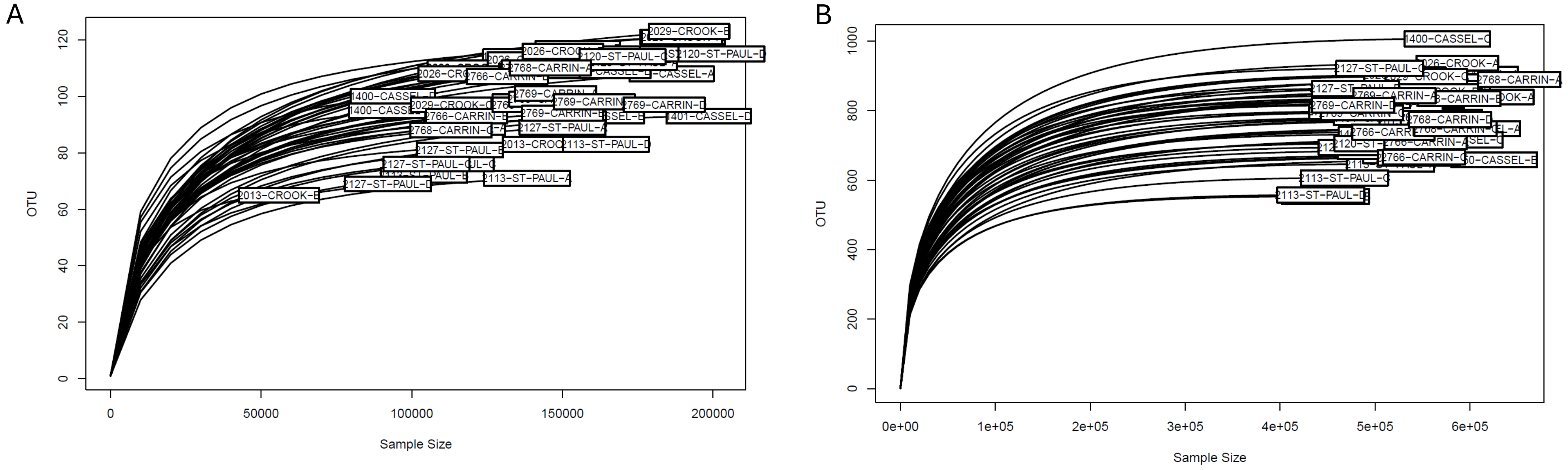


**S1 Fig. S1. Rarefaction curves for malting barley seed endophytes across four locations**. Rarefaction curves were made for all samples to evaluate the species richness, depth of sampling and sequencing coverage for bacterial 16S (A) and fungal ITS (B).

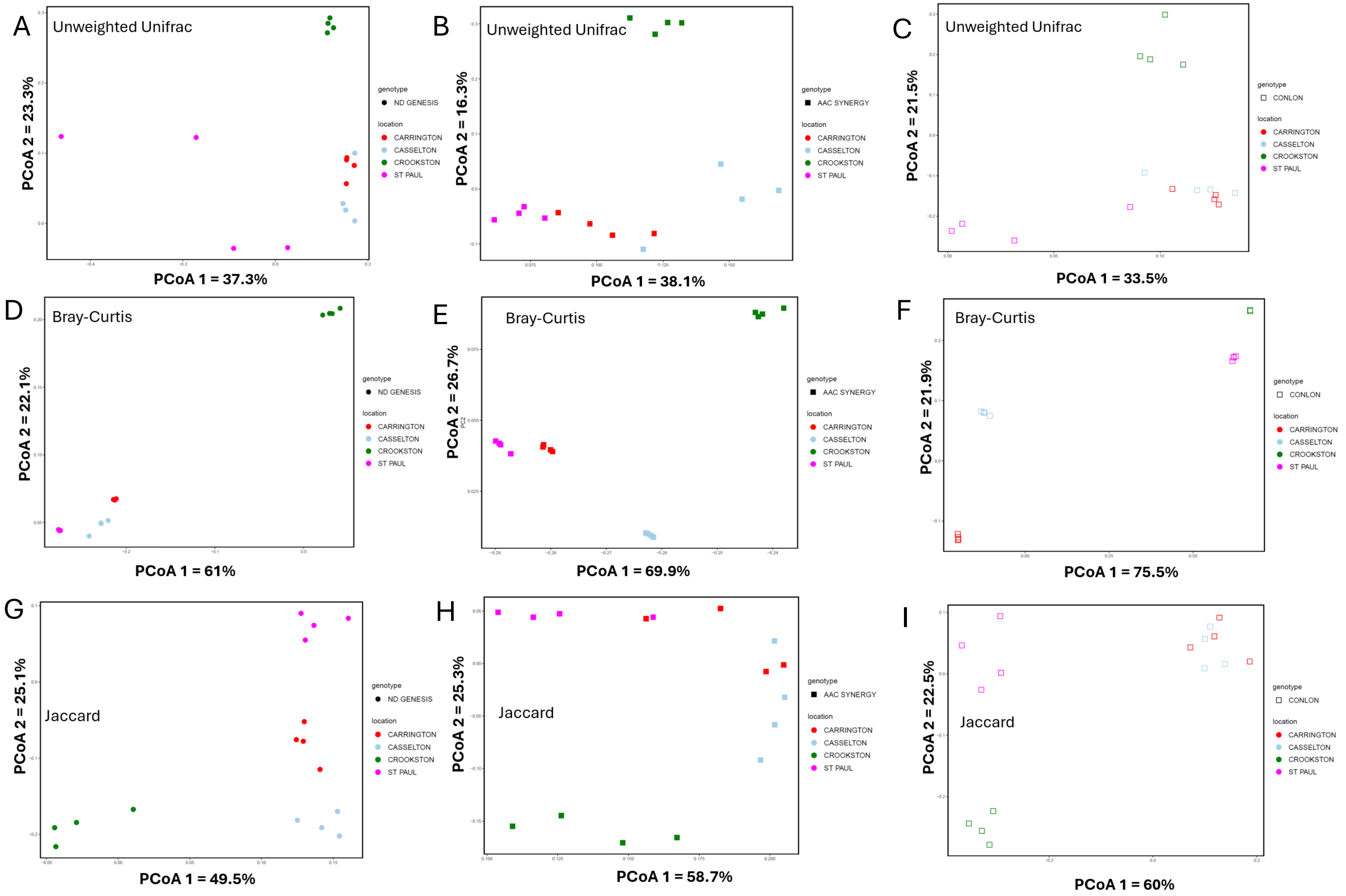


**S1 Fig. S2. - Bacterial community composition of malting barley seed endophytes across four locations.** A-I represents the PCoA of 16S rRNA amplicon sequencing data across location for each cultivar based on unweighted unifrac (A-C), Bray-Curtis (D-F) and Jaccard distances (G-I). A,E,H are for ND Genesis; B,E,H are for AAC Synergy while C,E,H are for Conlon genotypes.

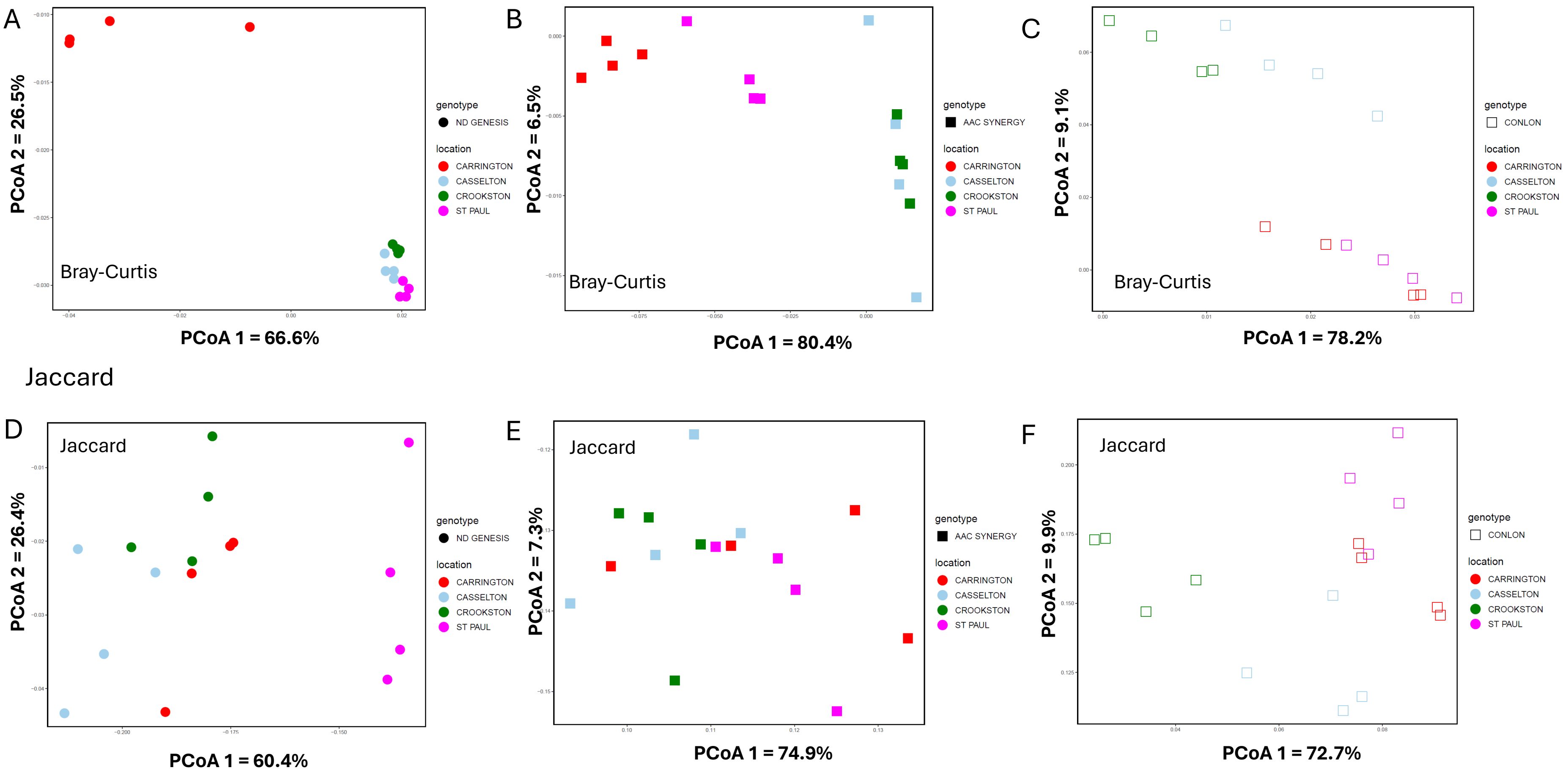


**S1 Fig. S3. Fungal community composition of malting barley seed endophytes across four locations**. A-F represents the PCoA of fungal ITS sequencing data across location for each cultivar based on Bray-Curtis (A-C) and Jaccard distances (D-F). A,D are for ND Genesis; B, E are for AAC Synergy and C,F are for Conlon genotypes/cultivars.

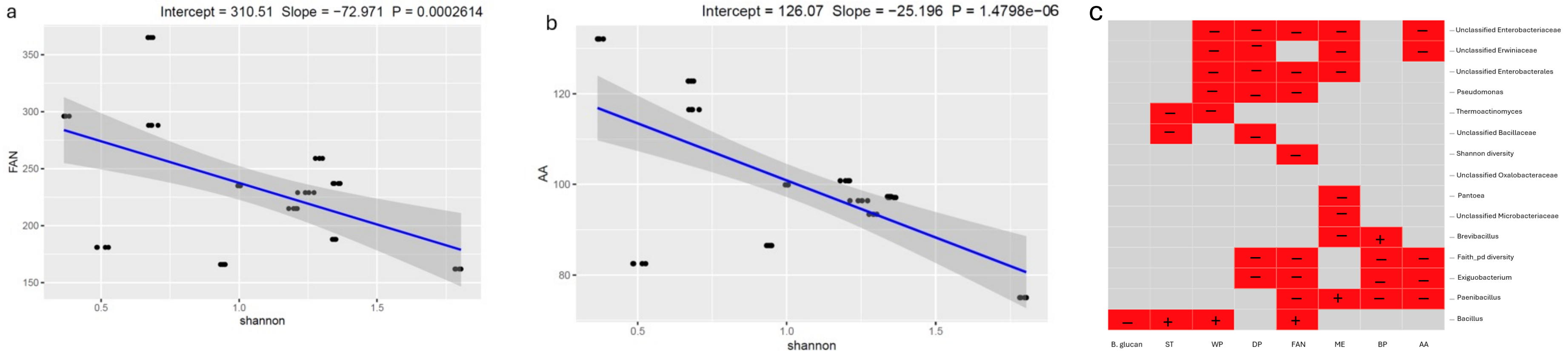


**S1 Fig. S4. Negative relationship between Shannon index, bacterial genus taxa and some malt quality traits**. For free amino nitrogen (FAN) (A), and alpha amylase (AA) (**B**) based on linear regression after adjusting for covariates genotype and location. (C) Association analyses between microbial taxa (genera) and malt quality traits using linear mixed model after adjusting for genotype and location effects. Only genera significantly associated with malt quality traits were shown (except for unclassified Oxalobacteraceae which was included to illustrate no significance with any malt quality trait) . + indicates positively association; - indicates negative association and red indicates significant.

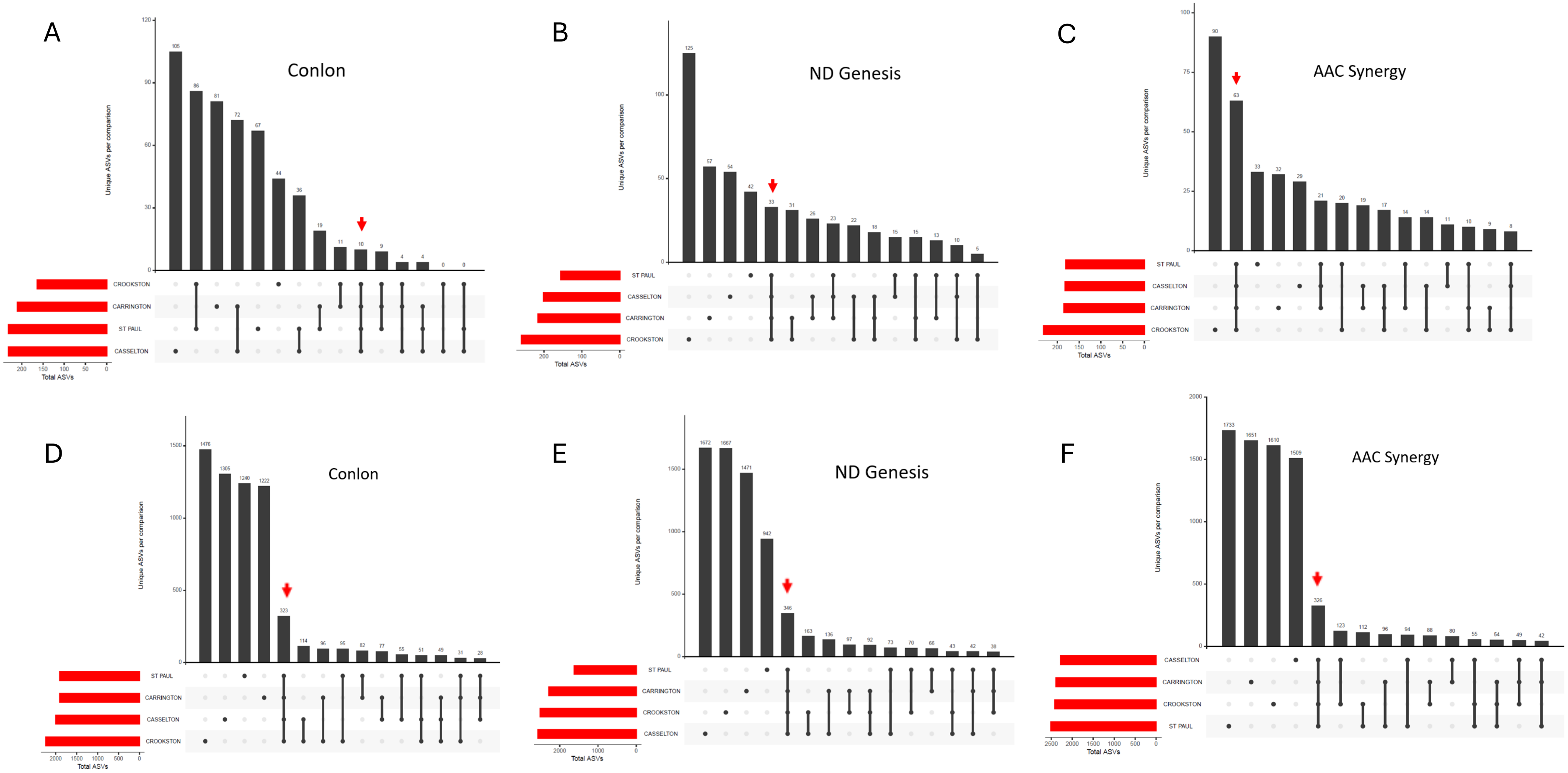


**S1 Fig. S5. UpSet plots of bacterial and fungal ASVs present in each location of Conlon, ND Genesis and AAC Synergy.** (A-C) UpSet plot showing the number of bacterial ASVs that are shared between or are unique for all locations for Conlon (A), ND Genesis (B) and AAC Synergy (C). (D-F) UpSet plot showing the number of fungal ASVs that are shared between or are unique for all locations for Conlon (D), ND Genesis (E) and AAC Synergy (F). Red arrow in A-F indicates shared ASVs common to all locations, while filled-in black dots with an edge between the dots indicates that these ASVs are present in multiple locations.

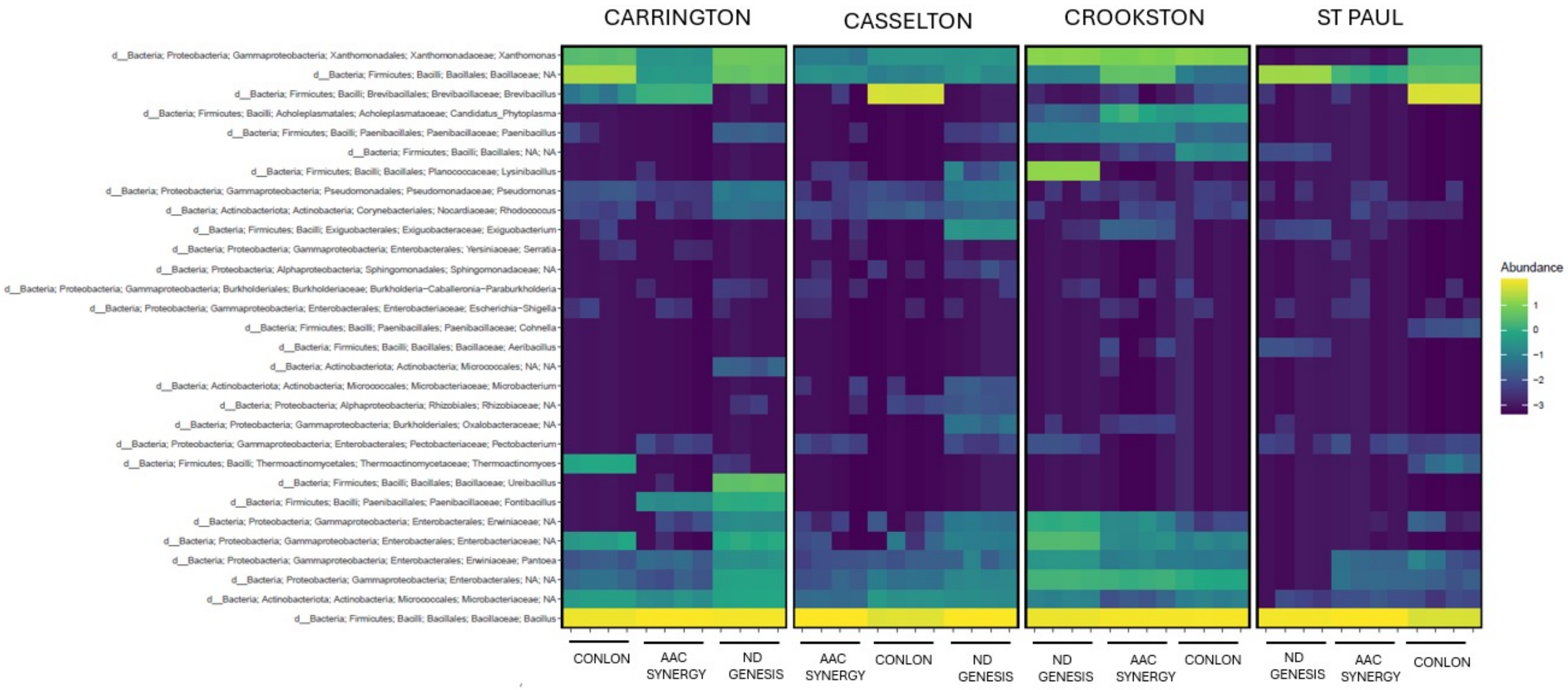


**S1 Fig. S6. Heatmap of the relative abundance of the top 30 bacterial seed endophytic communities of malting barley genotypes/cultivars across four locations.**

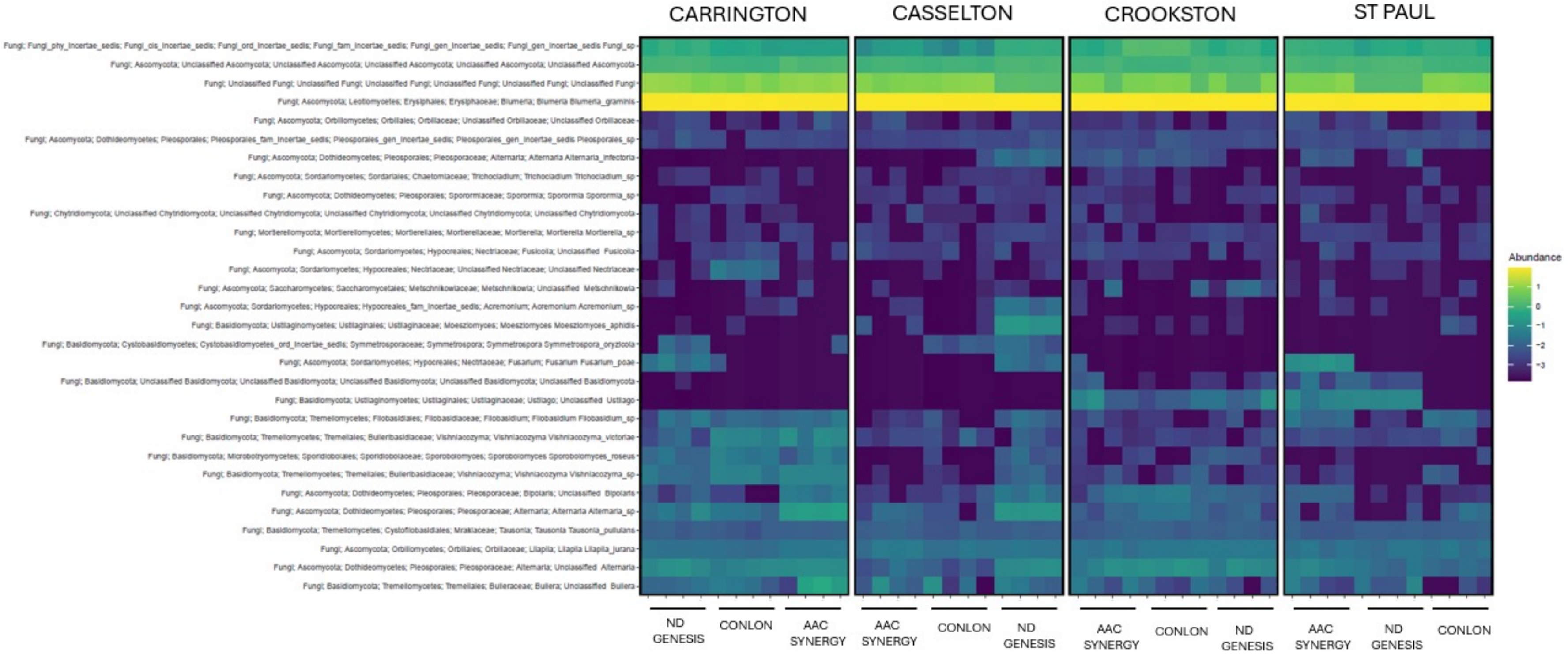


**S1 Fig. S7. Heatmap of the relative abundance of the top 30 fungal seed endophytic communities of malting barley genotypes/cultivars across four locations.**

A

| Feature ID | Sequence Length | Sequence |
|---|---|---|
| 7fc1d6e92ee03a814aa144f32e1f63d0 | 260 | CCTACGGGAGGCAGCAGTAGGGAATCTTCCGCAATGGACGAAAGTCTGACGGAGCAACGCCGCGTGAGTGATGAAGGTTTTCGGATCGTAAAACTCTGTT |
| eead808e0197137883dc138bb21a8bdc | 260 | CCTACGGGAGGCAGCAGTGGGGAATTTTCCGCAATGGGCGAAAGCCTGACGGAGCAATGCCGCGTGGAGGTGGAAGGCCTACGGGTCGTCAACTTCTTTT |
| 93c2bc2e9cbd4f85f384a1d7eb5f5377 | 260 | CCTACGGGGGGCAGCAGTGGGGAATCTTGGACAATGGGCGAAAGCCCGATCCAGCAATATCGCGTGAGTGAAGAAGGGCAATGCCGCTTGTAAAGCTCTT |
| 293321095e4cf6dfe46bedf510de6134 | 260 | CCTACGGGAGGCAGCAGTAGGGAATCTTCCGCAATGGACGAAAGTCTGACGGAGCAACGCCGCGTGAGTGATGAAGGTTTTCGGATCGTAAAACTCTGTT |
| 0810b374c63fb741fcee68eb79045c46 | 260 | CCTACGGGAGGCAGCAGTAGGGAATCTTCCGCAATGGACGAAAGTCTGACGGAGCAACGCCGCGTGAGTGATGAAGGTTTTCGGATCGTAAAACTCTGTT |
| 808829447d35304c65d40ab5807f4325 | 260 | CCTACGGGAGGCAGCAGTAGGGAATCTTCCGCAATGGACGAAAGTCTGACGGAGCAACGCCGCGTGAGTGATGAAGGTTTTCGGATCGTAAAGCTCTGTT |
| 9bc848531af1ba122a1e1e28941b935f | 260 | CCTACGGGTCGTCAACTTCTTTTCTCGGAGAAGAAACAATGACGGTATCTGAGGAATAAGCATCGGCTAACTCTGTGCCAGCAGCCGCGGTAAGACAGAG |
| e28abed63dd9831365aaced43cd1f049 | 260 | CCTACGGGAGGCAGCAGTGGGGAATATTGGACAATGGGCGCAAGCCTGATCCAGCCATGCCGCGTGGGTGAAGAAGGCCTTCGGGTTGTAAAGCCCTTTT |

**Sequences producing significant alignments:**

Select: All None Selected:0

Alignments Download GenBank Graphics Distance tree of results

| | Description | Max Score | Total Score | Query Cover | E value | Per. Ident | Accession |
|---|---|---|---|---|---|---|---|
| ☐ | Xanthomonas translucens strain NRCIB_X6 16S ribosomal RNA gene, partial sequence | 470 | 470 | 100% | 1e-127 | 100.00% | gi\|2148246466\|OL504772.1 |

B

Download GenBank Graphics

Xanthomonas translucens strain NRCIB_X6 16S ribosomal RNA gene, partial sequence

**Sequence ID:** gi|2148246466|OL504772.1 **Length:** 1391 **Number of Matches:** 1

**Range 1: 285 to 544** GenBank Graphics ▼ Next Match ▲ Previous Match

| Score | Expect | Identities | Gaps | Strand |
|---|---|---|---|---|
| 470 bits(520) | 1e-127 | 260/260(100%) | 0/260(0%) | Plus/Plus |

```
Query  1    CCTACGGGAGGCAGCAGTGGGGAATATTGGACAATGGGCGCAAGCCTGATCCAGCCATGC  60
            ||||||||||||||||||||||||||||||||||||||||||||||||||||||||||||
Sbjct  285  CCTACGGGAGGCAGCAGTGGGGAATATTGGACAATGGGCGCAAGCCTGATCCAGCCATGC  344

Query  61   CGCGTGGGTGAAGAAGGCCTTCGGGTTGTAAAGCCCTTTTGTTGGGAAAGAAAAGCAGTC  120
            ||||||||||||||||||||||||||||||||||||||||||||||||||||||||||||
Sbjct  345  CGCGTGGGTGAAGAAGGCCTTCGGGTTGTAAAGCCCTTTTGTTGGGAAAGAAAAGCAGTC  404

Query  121  GGTTAATACCCGATTGTTCTGACGGTACCCAAAGAATAAGCACCGGCTAACTTCGTGCCA  180
            ||||||||||||||||||||||||||||||||||||||||||||||||||||||||||||
Sbjct  405  GGTTAATACCCGATTGTTCTGACGGTACCCAAAGAATAAGCACCGGCTAACTTCGTGCCA  464

Query  181  GCAGCCGCGGTAATACGAAGGGTGCAAGCGTTACTCGGAATTACTGGGCGTAAAGCGTGC  240
            ||||||||||||||||||||||||||||||||||||||||||||||||||||||||||||
Sbjct  465  GCAGCCGCGGTAATACGAAGGGTGCAAGCGTTACTCGGAATTACTGGGCGTAAAGCGTGC  524

Query  241  GTAGGTGGTTGTTTAAGTCC  260
            ||||||||||||||||||||
Sbjct  525  GTAGGTGGTTGTTTAAGTCC  544
```

**S1 Fig. S8. Feature ID sequence identification and BlastN analysis of 16S rRNA partial sequence of Xanthomonas.** (A) Feature ID colored in orange represents genus Xanthomonas (B) BlastN analysis of the 260 bp sequence shared 100% homology with Xanthomonas translucens.

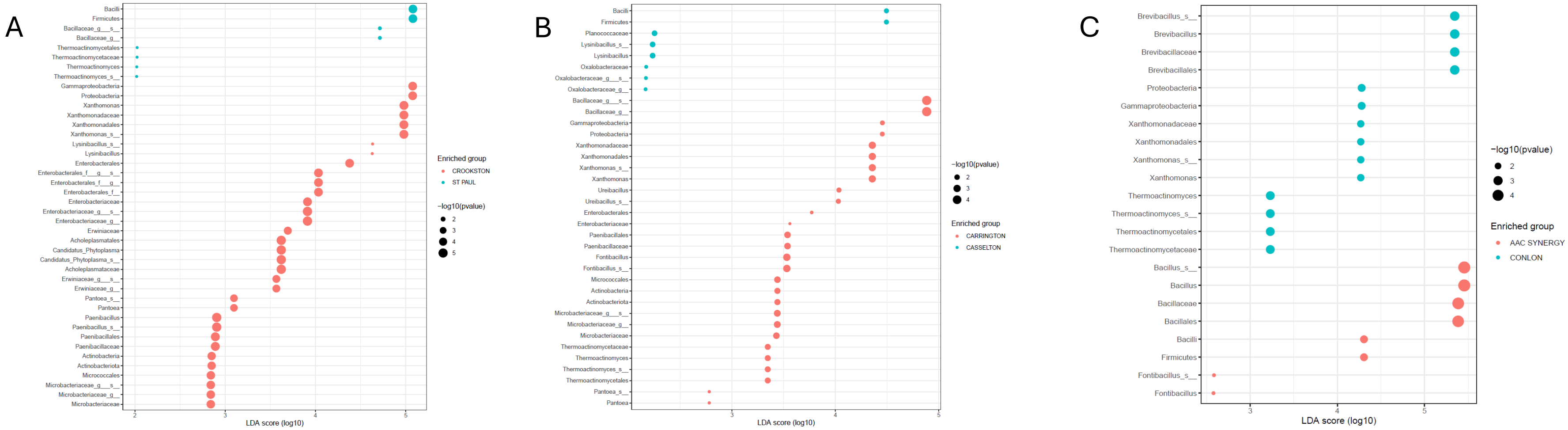


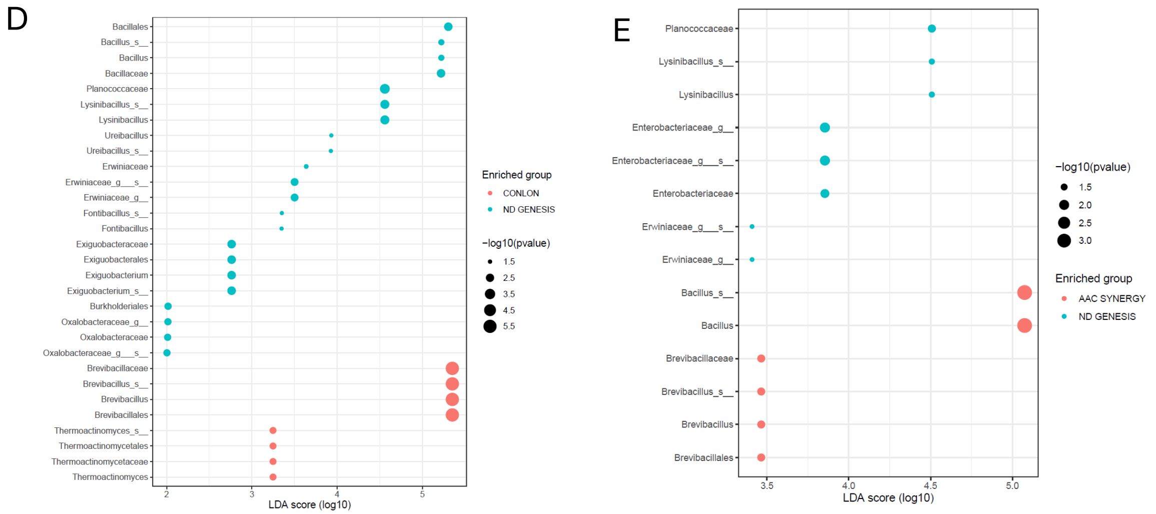


S1 Fig. S9. **Genotype and location pairwise comparison of the seed endophytic bacteria microbial taxonomic composition.** For Crookston vs. St. Paul (A), Carrington vs. Casselton (B), AAC Synergy vs. Conlon (C), Conlon vs. ND Genesis (D), and AAC Synergy and ND Genesis (E). For A-E, dot plots of the LDA scores (log 10) computed for features with differential abundance in each pairwise assessments as indicated above.

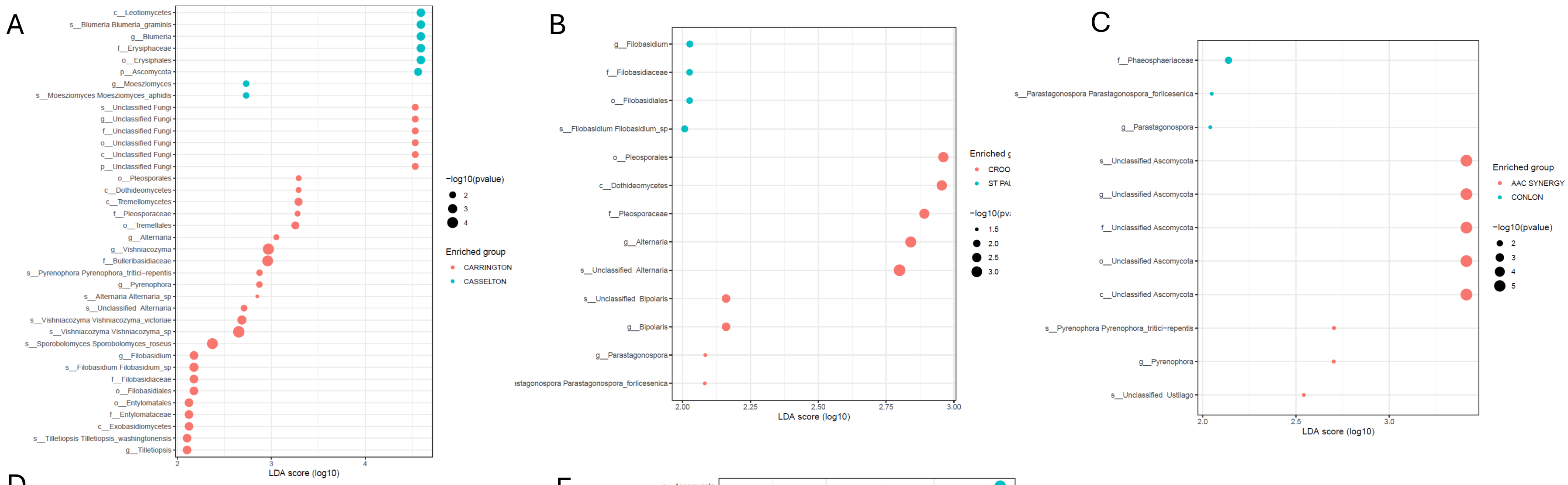

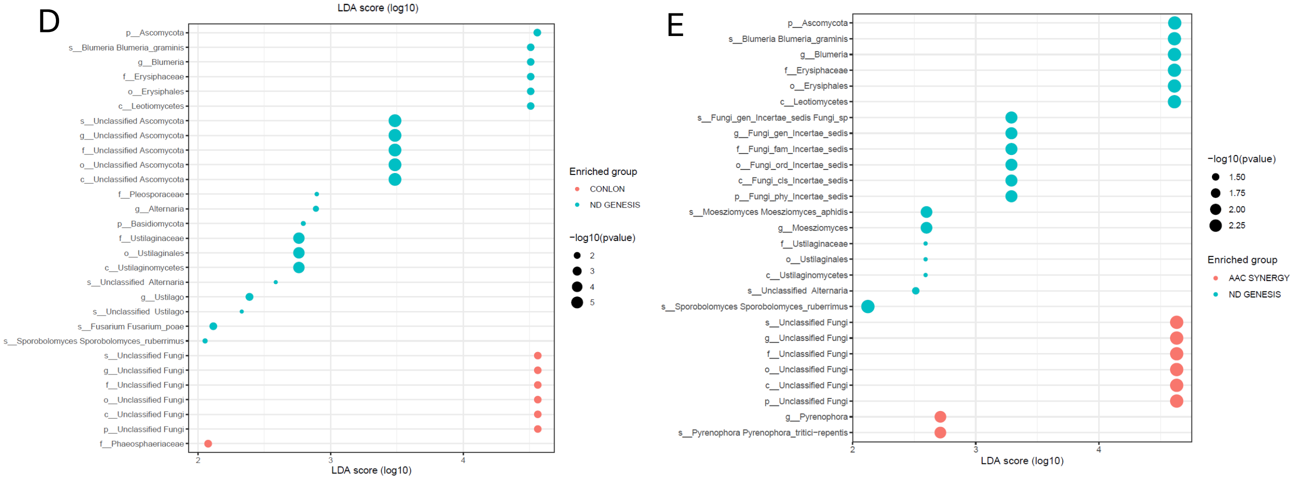


**S1 Fig. S10. Genotype and location pairwise comparison of the seed endophytic fungal microbial taxonomic composition.** For Carrington vs. Casselton (A), Crookston vs. St. Paul (B), AAC Synergy vs. Conlon (C), Conlon vs. ND Genesis (D), and AAC Synergy and ND Genesis (E). For A-E, dot plots of the LDA scores (log 10) computed for features with differential abundance in each pairwise assessments as indicated above.
